# Inherent protection of bacteria from beta-lactam antibiotics by wet-dry cycles with microscopic surface wetness

**DOI:** 10.1101/2020.11.09.373787

**Authors:** Yana Beizman-Magen, Maor Grinberg, Tomer Orevi, Nadav Kashtan

**Affiliations:** Department of Plant Pathology and Microbiology, Institute of Environmental Sciences, Robert H. Smith Faculty of Agriculture, Food, and Environment, Hebrew University, Rehovot, 76100 Israel

## Abstract

A large portion of bacterial life occurs on surfaces that are not constantly saturated with water and experience recurrent wet-dry cycles. While soil, plant leaves and roots, and many indoor surfaces may appear dry when not saturated with water, they are in fact often covered by thin liquid films and microdroplets, invisible to the naked eye, known as microscopic surface wetness (MSW). Such MSW, resulting from the condensation of water vapor to hygroscopic salts, is ubiquitous yet largely underexplored. A wide variety of antibiotics are abundant in environments where MSW occurs, yet little is known about bacterial response to antibiotics in wet-dry cycles and under MSW conditions. Using *E. coli* as a model organism, we show, through a combination of experiments and computational modeling, that bacteria are considerably more protected from beta-lactams under wet-dry cycles with MSW phases, than they are under constantly wet conditions. This is due to the combined effect of several mechanisms, including tolerance triggered by inherent properties of MSW, i.e., high salt concentrations and slow cell growth, and the deactivation of antibiotics due to physicochemical properties of MSW. Remarkably, we also find evidence for a cross-protection effect, where addition of lethal doses of antibiotic before drying significantly increases cells’ survival under MSW. As wet-dry cycles with MSW and beta-lactams, as well as other antibiotics, are common in vast terrestrial microbial habitats, our findings are expected to have significant implications for how we understand antibiotic response, population dynamics, and interspecies interactions in these globally important microbial ecosystems.

## Introduction

Bacteria are often found on natural and artificial surfaces that are not constantly wet, and experience frequent drying and rewetting. During ‘dry’ periods, though many of these surfaces appear dry to the naked eye, they are typically covered by thin liquid films and micrometre-sized droplets, termed microscopic surface wetness (1, 2) (MSW; Fig. 1). Key to the formation and retention of microscopic wetness is the presence of deliquescent substances – mostly highly hygroscopic salts – that absorb moisture from the air until they dissolve-in and form a liquid solution (3, 4). Thus, residual deposits of deliquescent compounds that cover a surface, become, or are retained as, MSW when the relative humidity (RH) exceeds their specific deliquescence, or efflorescence, points. As deliquescent substances are ubiquitous, MSW occurs in many microbial habitats that undergo wet-dry cycles, including plant root and leaf surfaces (1, 5, 6), soil and rock surfaces (7, 8), the built environment (9, 10), and probably even on human and animal skin (11) (Fig. 1). Hence, microscopic surface wetness is a common hydration condition in diverse important microbial habitats.

**Fig. 1.**
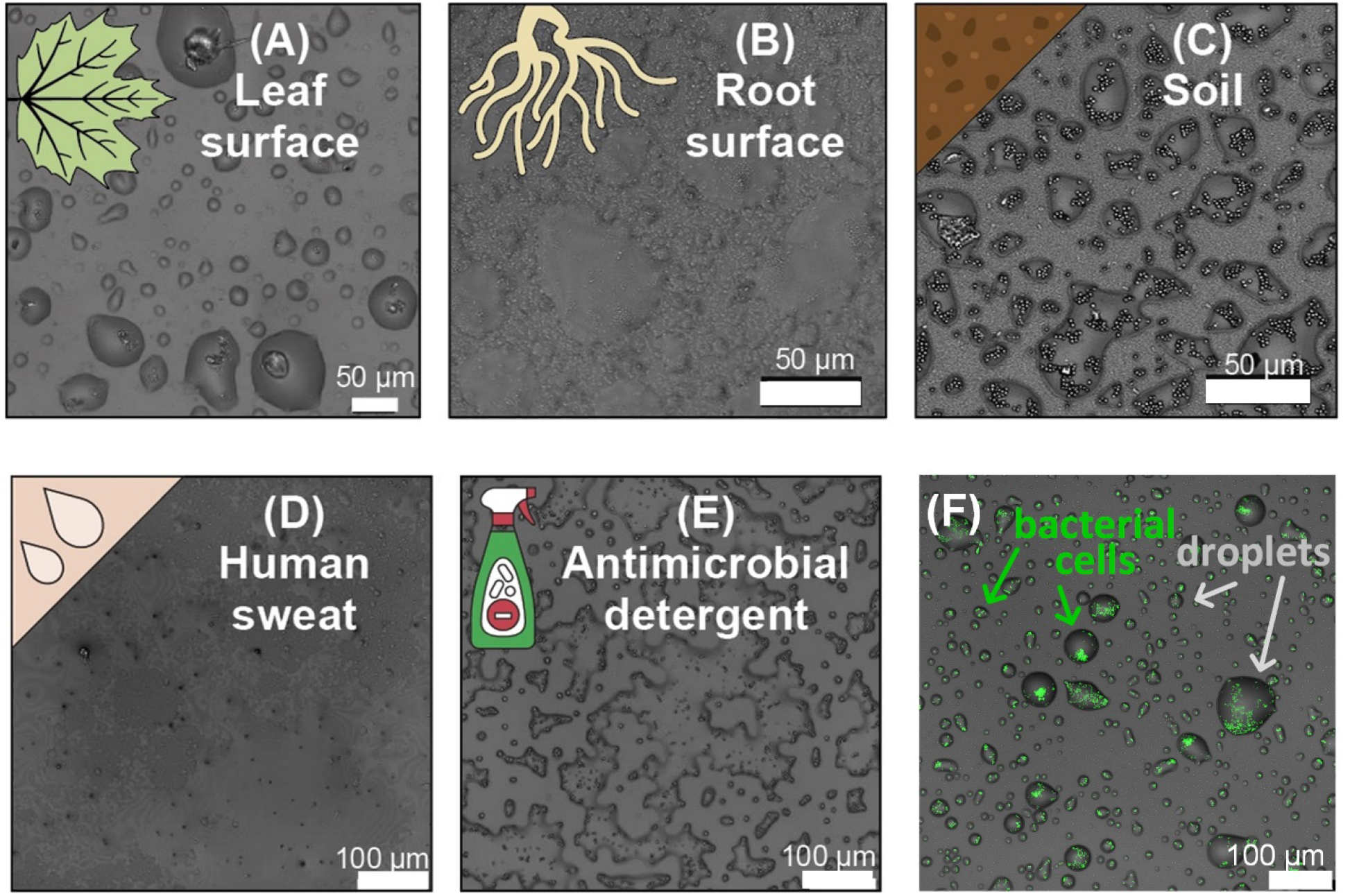
Microscopic surface wetness (MSW) occurs in many ubiquitous natural and artificial surfaces. MSW from natural samples dried on glass surfaces at 60-70% RH. **A.** Leaf wash. **B.** Root wash. **C.** Soil extraction. **D.** Human sweat (representing the skin environment). **E.** Indoor MSW resulting from the use of antimicrobial detergent. **F.** Fluorescently-labeled bacteria in microscopic surface wetness (M9 medium): Cells reside within stable micro-meter-sized droplets. Solutions in droplets are highly concentrated (high salinity); cells are stressed, and growth rates are low.

MSW can protect bacteria from complete desiccation (2). For example, MSW resulting from the deliquescence of hygroscopic aerosols – which are ubiquitous on leaves (12, 13) – may explain how bacteria survive daytime dryness on plant leaves (2, 14). However, while possibly protecting from desiccation, MSW is a harsh micro-environment that differs from bulk liquid. MSW’s physicochemical properties such as extremely high salinities, altered pH, increase in reactive oxygen species (ROS), and segregation into tiny droplets (15–17), necessarily impose severe stresses on cells are likely to significantly impact the physiology of bacterial cells therein (18–20). Cells that are exposed to drying and MSW were shown to activate stress responses (21) and exhibit very low, or even close to zero, growth rates (2).

A large variety of antimicrobial compounds are abundant in environments where wet-dry cycles and MSW occur. For example in soil, leaf, and root surfaces, major sources of these antimicrobials are their natural production by microbes and plants (22–25), and their release into water and soil by human activity and agricultural practices (26–28). These antibiotic compounds produced and secreted by microbes are the driving force of microbial warfare and competition, and are consequently expected to impact population dynamics and community compositions of natural microbial consortia. Little is known, though, about bacterial response to antibiotics on surfaces with frequent wet-dry cycles and under MSW conditions.

To begin to understand antibiotic response under wet-dry cycles and in MSW, we chose to focus on beta-lactams – the oldest and one of the most commonly used antibiotics (29–33). Beta-lactams target penicillin-binding-proteins in bacterial cell membranes that are responsible for crosslinking peptidoglycan layers, resulting in accumulated damage to cell membrane that eventually leads to cell lysis. In addition to their clinical importance, diverse groups of beta-lactams are abundant in natural environments, where they are produced by many bacteria and fungi (25, 31).

Bacteria respond to the presence of antibiotics in various ways. Resistance to antibiotics is the (usually inherited) ability of bacteria to grow under high antibiotic concentrations, i.e., above the minimum inhibitory concentration (MIC) (34, 35). Tolerance is the ability of a microorganism to survive a transient exposure to an antibiotic at concentrations that would otherwise be lethal (i.e., above the MIC) (34, 36, 37). In many cases, tolerance can be thought as a transient protection that emerges under environmental conditions that impose low metabolic activity and reduced growth, may result from collective properties of populations rather than of individual cells, or are triggered by environmental stressors, which in turn confer lower susceptibility to an antibiotic (34, 37–40).

Two key properties of MSW as a microbial environment – slow bacterial growth rates, and high salt concentrations – have been shown to increase tolerance and reduce susceptibility to antibiotics (41–47). As beta-lactam antibiotics inhibit proper cell wall synthesis, faster growing cells accumulate more damage, leading to a higher likelihood of death (41–43). Indeed, robust correlations were found between growth and lysis rates in diverse pairs of beta-lactams and bacteria (41, 47). Another inherent feature of MSW is high salinity, which results from the high concentrations of solutes (e.g., deliquescent salts) due to water evaporation. High salt concentrations were shown to reduce susceptibility to antibiotics through a cross-protection effect that results from the increased expression of efflux pumps (45, 46). Finally, the high salinity, ROS, and altered pH in MSW might affect the stability and activity of many antibiotic compounds, including beta-lactams (48).

We thus hypothesize that under wet-dry cycles with an MSW phase, bacteria are less susceptible to antibiotics in comparison to constantly wet (water-saturated) conditions. To test this hypothesis, we quantitatively studied *E. coli* populations’ responses to two beta-lactams: Ampicillin, and Carbenicillin (which is a more stable beta-lactam).

In this work, we build upon an experimental system developed in our lab that enables us to culture bacteria in wet-dry cycles with MSW phase, and to quantitatively estimate killing rates and cell survival based on Live/Dead assays and direct CFU counts. We compared survival of *E. coli* exposed to differing doses of two common beta-lactam antibiotics in wet-dry cycles with its survival under constantly wet conditions. Then we combined computational models with our experimental results to elucidate the mechanisms underlying the observed antibiotic response, and to further develop mechanistic understanding of antibiotic response under wet-dry cycles with MSW.

## Results

### Experimental system to study bacterial response to antibiotics under wet-dry cycles with MSW phase

To assess survival of *E. coli* Dh5α in response to exposure to beta-lactams, we built upon a microscopy-based experimental system that enabled us to study bacteria under varying hydration conditions and MSW (2). Our wet-dry cycle experiments consisted of three sequential phases: drying, MSW, and rewetting (Supp. Fig. S1). Briefly, bacterial cells were loaded into 1 μl drops of M9 minimal media supplemented with ampicillin (Amp) or carbenicillin (Crb) in concentrations of 5 μg/ml (2X MIC), 50 μg/ml (20X MIC), or without antibiotics. The drops were incubated in an on-stage environmental chamber under constant temperature and RH (32°C; 70% RH). In about two hours, the drops dried out, and MSW conditions – in the form of stable thin liquid films – were established. MSW conditions, which constituted the ‘dry period’, were maintained for either one hour (short cycle) or 12 hours (long cycle) before rewetting. Rewetting was executed by increasing RH followed by the addition of broth medium (see Methods, Fig. 2A, Supp. Fig. S1). Corresponding experiments in constantly wet conditions, with equivalent initial cell density, media, and antibiotic concentrations, were conducted likewise (representing continuous water-saturated conditions - see Methods).

**Fig. 2.**
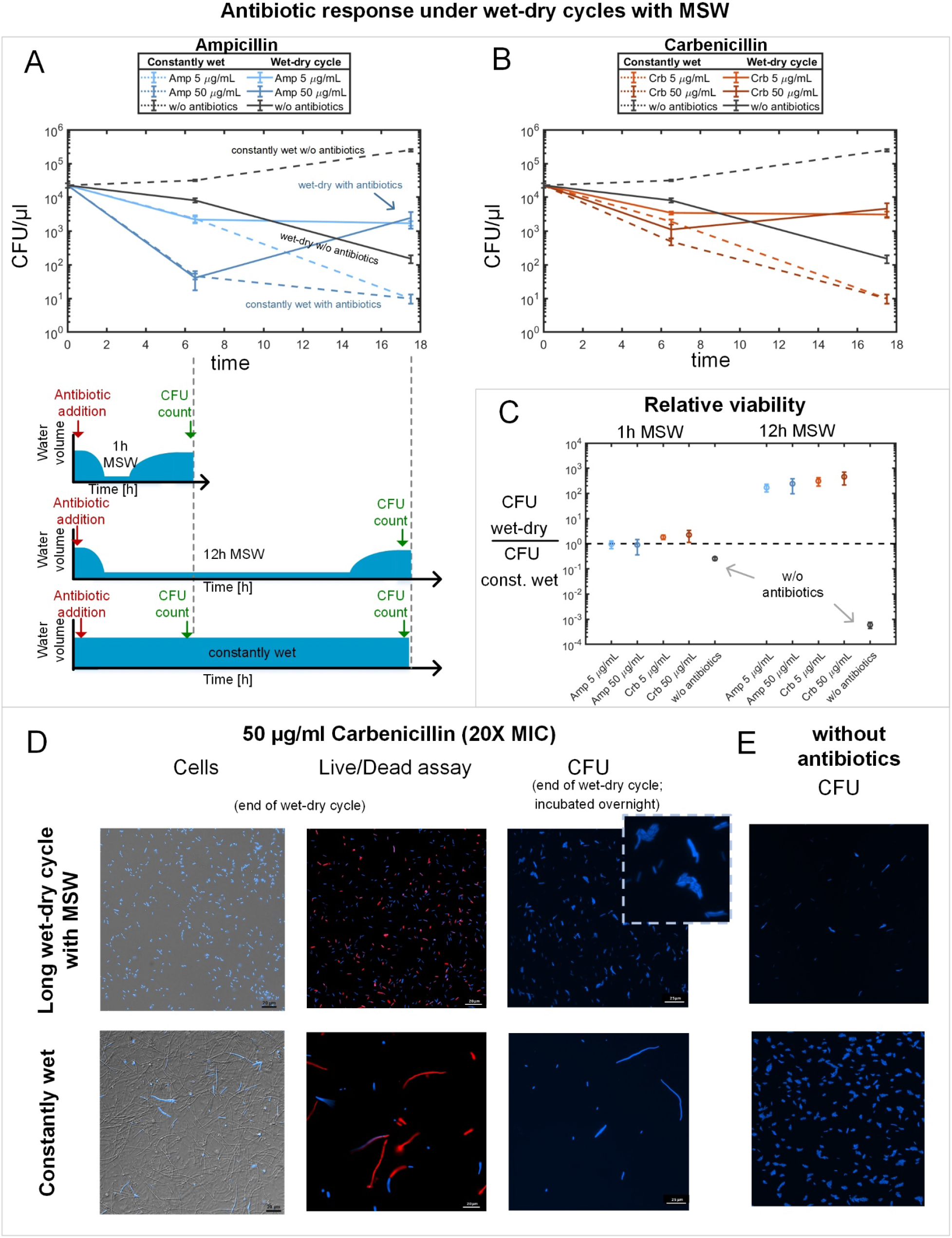
Wet-dry cycles with MSW protect bacteria from beta-lactam antibiotics. **A.** Bacterial response to Amp under short and long drying-rewetting cycle, and under constantly wet conditions. Mean ± SE CFU/μl are at times 0, 6.5h, and 17.5h. Amp was added at concentrations of 2X MIC and 20X MIC; no antibiotic was added in controls. CFU assay was conducted as described in Methods. **B.** same as (A) but with Crb. **C.** Relative viability (Mean ± SE) at the end of a wet-dry cycle compared to that under constantly wet conditions, based on CFU counts. **D.** Microscopy images of live cells (blue), Live/dead assay (live in blue, dead in red), and micro-colonies formed on agarose pads (see Methods) placed at the end of the wet-dry cycle with 20x MIC Crb, compared to constantly wet conditions with same Crb Conc. **E.** Micro-colonies formed from the end of an experiment without antibiotics (t = 17.5h).

To estimate bacterial survival, we developed a method of counting micro-colonies directly in our experimental system: This was done by laying agarose pads on the colonized surface at the end of the rewetting phase and then, following overnight growth, scanning the surface under the microscope to count micro-colonies (CFU/μl counts - see details in Methods). As a complementary method for measuring cell survival, we used propidium iodide staining to distinguish between live and dead cells during the rewetting phase (live/dead assays - see Methods).

### Inherent protection from beta-lactams under wet-dry cycle with MSW phase

Under constantly wet (water-saturated) conditions, there was a rapid killing of most bacteria – consistent with previously reported time kill curves (49). At 6.5h, the equivalent time of the short cycle experiment, there was about 10-fold reduction in CFU/μl counts for the low antibiotic concentration (2X MIC; less than 10% of the initial cell population were viable) and ~100 to 1,000-fold reduction of CFU/μl counts for the high antibiotic concentration (20X MIC; 0.1% to 1% of cells remain viable) (Fig. 2 A-B). At 17.5h, the equivalent time of the long cycle experiment, there were < 10 CFU/μl, representing < 0.1% viable cells of the initial cell population, for both antibiotics types and concentrations (Fig. 2 A-B).

Remarkably, in the long wet-dry cycle experiments where cells were exposed to 12 h of MSW, there were >1,000 CFU/μl (Fig. 2 A-B), representing at least two orders of magnitude higher CFU than under constantly wet conditions, at the corresponding time (t = 17.5 h) (Fig. 2 C). In the short wet-dry cycle experiments, however, we saw either similar CFU counts as under constantly wet conditions at the corresponding time (Amp experiments), or only modest increase in CFU counts (Crb experiment) (Fig 2A-C). Similar trends, though with an order of magnitude smaller differences, were observed by live/dead assays for cell survival rather than CFU counts (Supp. Fig. S2, S3). In the long wet-dry cycle experiments, there were about 10 times more live cells than in the corresponding constantly wet experiments (Supp. Fig. S3), as opposed to 100 times more viable cells based on CFU counts (live cell count included intact cells expressing the transformed BFP2 gene and not stained by propidium iodide).

Inspection of the microscopy images confirmed that the number of cells under constantly wet conditions with antibiotics was indeed much smaller than in the equivalent wet-dry cycle experiment (Fig. 2D, left column, Supp. Fig. S4), likely because many cells were lysed due to cell wall deterioration and osmotic rupture (50). These lysed cells did not even appear stained in red by propidium iodide. Many of the live cells that did survive were longer and thicker, a known effect of beta-lactams (Fig. 2D, Supp. Fig. S4, S5) (51). In contrast, under wet-dry cycles and MSW, while cells were not growing or dividing, most cells were intact, and there were many more live cells than there were under the corresponding constantly wet conditions (Fig. 2D, Supp. Fig. S4, S5). High correlation was found between the numbers of live cells and the number of CFUs (r = 0.998, P < 0.001, Supp. Fig. S6). The ratio of live to dead cells after 12h under MSW conditions was around 50% (Supp. Fig. S7), yet a comparison based on the percentage of dead cells would be misleading, as it does not count cells that were already severely ruptured (Fig. 2D, Supp. Fig. S4). Micro-colonies could be easily observed on the agarose pads that were placed on the well bottoms (see Methods), with significantly more colonies in the long wet-dry cycle experiments than under constantly wet conditions (Fig. 2 C,D, Supp. Fig. S8). Without antibiotics, there were indeed many more CFUs in the constantly wet experiment compared to the wet-dry cycle experiments (Fig. 2E, Supp. Fig. S8), reflecting high growth and high survival under wet conditions, and slow (or no) growth and lower survival in MSW.

### The underlying mechanisms of the observed protection from beta-lactams

To better understand the mechanisms underlying the observed protection, we examined several directions: deactivation of antibiotics under MSW conditions, tolerance by slow growth, and cross protection due to high salt concentrations.

We first sought to assess the stability of both antibiotics during the drying process and under MSW conditions, and test whether a decay in their activity can explain our results. We found that during drying followed by an MSW period, a portion of the antibiotics was deactivated (in particular Amp; less so Crb, Supp. Fig. S9, Supporting Information), likely via degradation – possibly due to the high salt concentrations (Supp. Fig. S10). The rates of antibiotic deactivation during drying and MSW were not sufficient, however, to explain the observed high cell survival at the end of the long wet-dry cycle, since high concentrations of active antibiotics were retained throughout the duration of the cycle. Moreover, as the concentration of all solutes increases during drying due to water evaporation, the antibiotic concentrations are expected to increase accordingly. In previous work, under similar conditions (medium and RH), we estimated a concentration factor of ~50 times (2). Thus, by the time MSW is formed, the antibiotic concentration in the experiment that began with 20X MIC is expected to reach ~1000X MIC (assuming most of the antibiotic is still active after 1h in MSW, Fig. S9). Even if we assume that only 10% of the antibiotic remains active at the end of a 12h MSW period, it is not clear how so many cells could survive a high antibiotic dosage of ~100X MIC. Thus, there must be additional mechanisms, other than decay of antibiotic activity, underlying the observed high bacterial survival.

Next, we sought to test the impact of salt concentrations on lysis rates. Cross protection due to high salt concentrations, which is one major stress associated with MSW, has been previously reported (46). During drying and formation of MSW, cells are exposed to rapid increase in salt concentration until stable concentrated solutions in the form of microdroplets or thin films are formed (with an estimated final concentration of about M9 ~25X (2)). Adopting a method suggested by Lee *et al.* (47), we estimated the lysis rates under a range of M9 medium concentrations. We found that at M9 concentrations ≥ 10X, lysis rates decreased significantly, approaching zero at M9 15X (Fig. 3). Lysis rates were also highly correlated to cell growth rates, under the tested M9 concentrations. At high salt concentrations, growth rates were low, as were estimated lysis rates (Fig. 3, Supp. Fig. S11), consistent with the well-known correlations between bacterial growth rates and beta-lactams’ response (41, 47). We therefore conclude that the high salinity leads to a pronounced reduction in lysis rates, via cross-protection or tolerance by slow growth, or both. Because high salinity is an inherent property of MSW, it likely plays a major role in the reduced lysis rates that we observed in the longer wet-dry cycle.

**Fig. 3.**
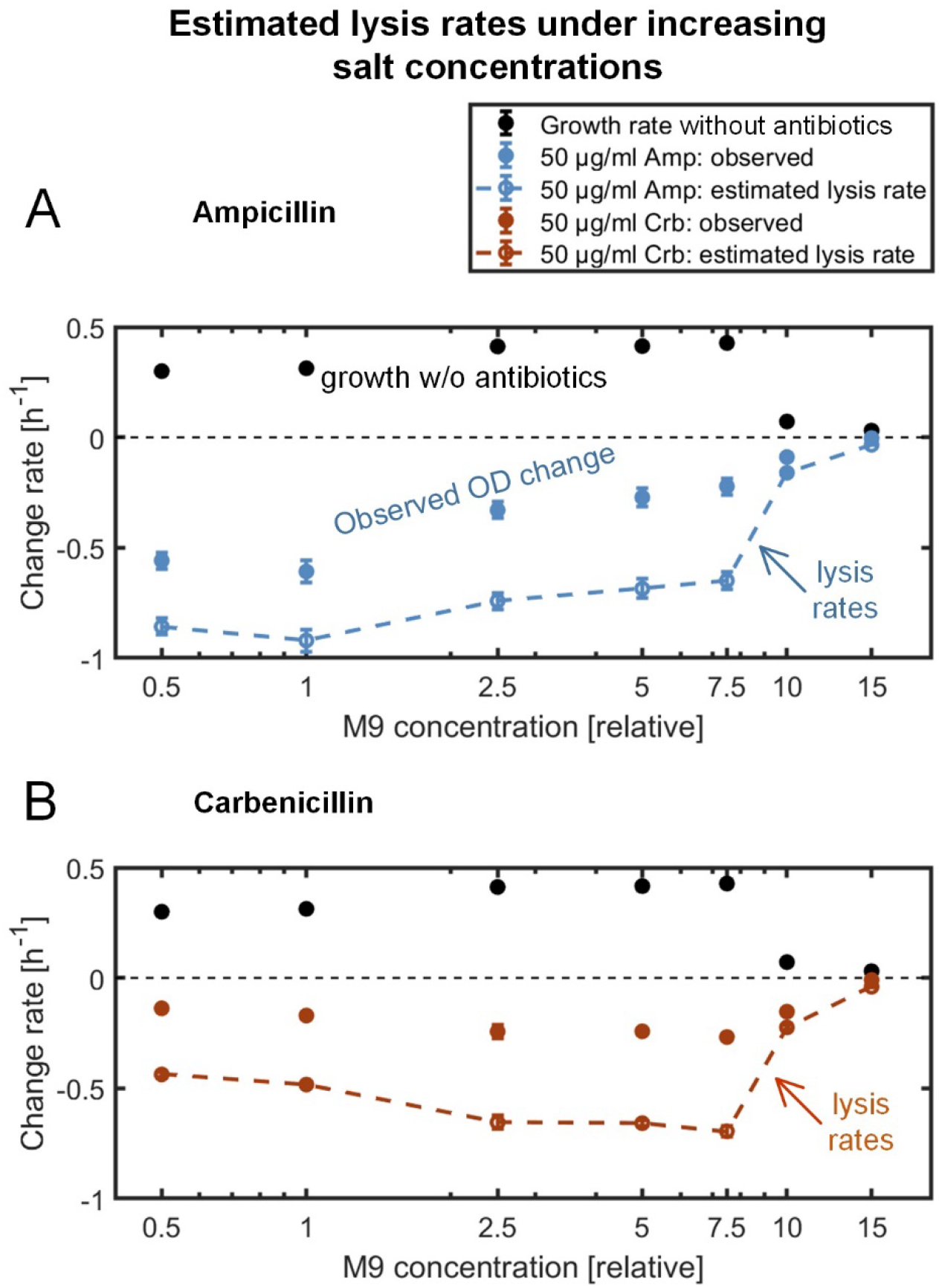
Estimation of growth and lysis rates in response to high concentration of antibiotics under increasingly concentrated M9 medium (see Methods). 50 μg/ml of beat-lactam antibiotic, Amp (**A**) or Crb (**B**) was added to a mid-exponential growth phase *E. coli* culture growing in various M9 concentrations. Lysis rates decrease significantly from M9 10X, approaching zero lysis at M9 15X.

### Beta-lactams cross-protect bacteria from drying and MSW conditions

Surprisingly, we observed that in the long wet/dry cycle experiment, cell survival was significantly higher in the experiments with antibiotics than they were in the corresponding experiments without antibiotics (Fig. 2 A-C, Supp. Fig. S3). That is, the addition of antibiotics before drying protected bacteria from stresses associated with drying and MSW conditions, and increased cell survival. Without antibiotics, about 1% of the cells were viable after 12h in MSW (CFU counts) (Fig. 2 A,B), while more than 10% were viable when antibiotics were added (Fig 2A,B). Increase in survival was also observed based on the live/dead assays, from 10% live cells without antibiotics to more than 40-50% live cells when antibiotic is added (Supp. Fig. S2, S3). This unexpected result was observed for both Amp and Crb, and at both tested concentrations (Fig. 2). These results point to an additional cross protection effect that has not been reported before, wherein antibiotics lead to cells’ increased protection from stresses associated with drying and MSW.

### A computational model provides a mechanistic understanding of experimental results

Finally, we aimed to develop a computational model that, based on our experimental results and their synthesis, can provide a mechanistic understanding of bacterial response to beta-lactams in wet/dry cycles with MSW phase. Toward this goal, we developed a mathematical model – a population-based ODE model – that incorporates the mode of action of the proposed mechanisms underlying increased bacterial protection from antibiotics under wet-dry cycles (see Methods and Supp. Info. ‘Model description’). Briefly, our model assumes that cell growth rates depend upon salt concentrations (i.e., medium conc.), and death rates depend upon antibiotic response, osmotic stress, and the observed reduced susceptibility to beta-lactams. The model takes into account the decay of antibiotic activity and the changes in water volume during drying and rewetting (Fig. 4A). Most model parameters were extracted from the experimental results (see Supp. Info ‘Model description’, Supp. Fig. S12, S13). The drying and rewetting dynamics, combined with antibiotic deactivation at high salt concentrations, can be seen as a trajectory in the ‘salt concentration x antibiotic concentration’ domain that dictates bacterial population dynamic (Fig. 4B). As can be seen in Fig. 4 D, E, this model produced results qualitatively similar to the experimental ones. The model suggests how changes in water levels, salt, and antibiotic concentrations, and the formation of MSW, govern the instantaneous death rate during the various phases (I, II, …, V) of a wet-dry cycle (Fig. 4A), in such a way that the resulting population sizes qualitatively agree with our experiments (Fig. 4 D,E, Supp. Fig. S14, S15). The model demonstrates the role that reciprocal cross-protection plays in the observed results, and how the duration of MSW conditions can affect the dynamics after rewetting, i.e., whether or not antibiotic concentration will inhibit growth (Fig. 4B). Importantly, incorporation of ‘tolerance by lag’ (34) after rewetting was required to achieve qualitative agreement with the experimental results. This lag phase, observed experimentally in delayed growth of cells after rewetting that correlates with the length of MSW conditions, was necessary to explain the reduced killing after rewetting in the long cycle. The model thus suggests that such tolerance ‘by lag’ likely plays a role in reduced overall bacterial susceptibility to beta-lactams, in particular when antibiotic concentrations are high and the ‘dry’ periods are long.

**Fig. 4.**
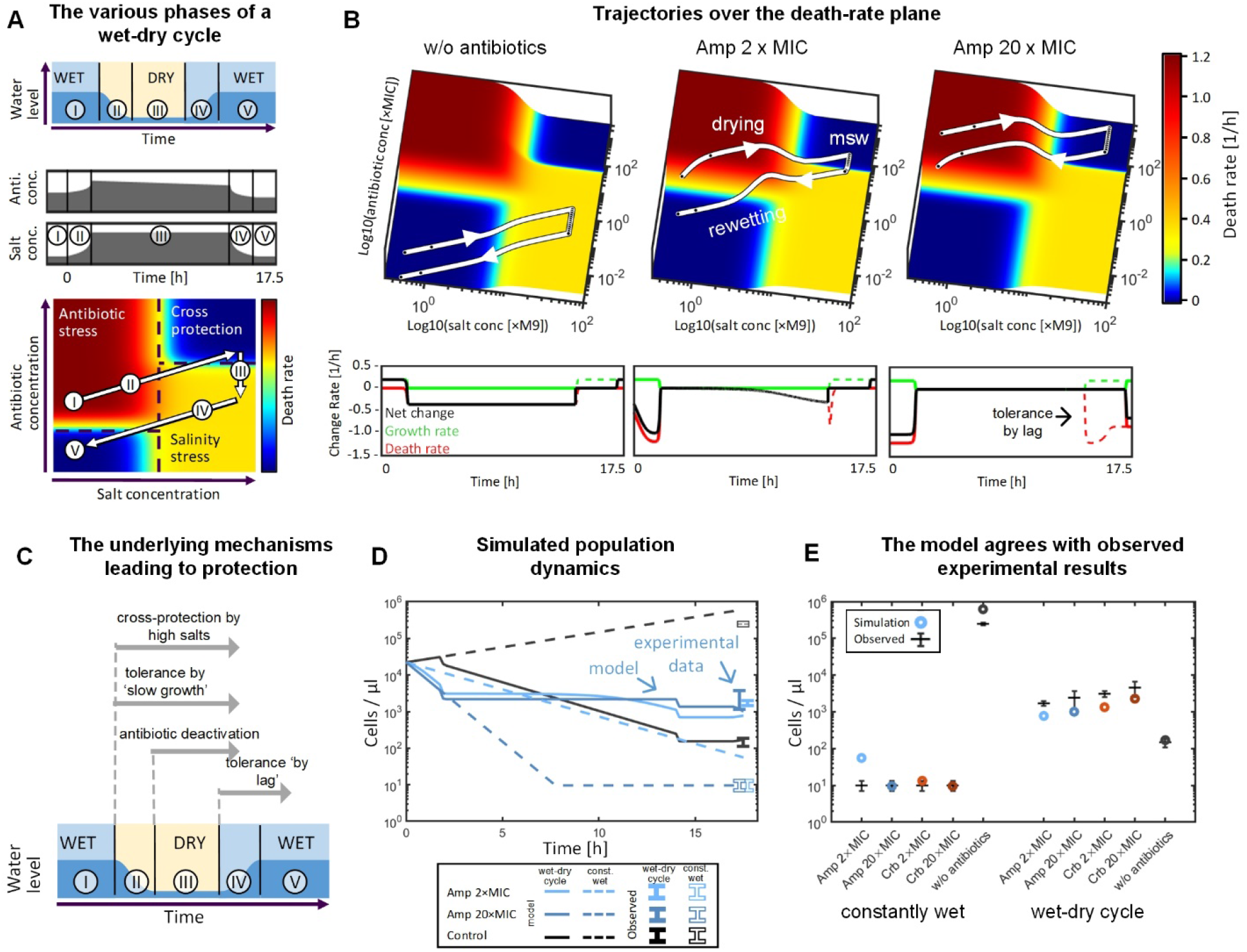
Mathematical modeling provides a mechanistic understanding of antibiotic response under wet-dry cycles with MSW. **A.** Under wet-dry cycle, varying water volume and antibiotic deactivation rates trace a trajectory on the salt conc. x antibiotic conc. 2D plane (white arrows). The bacterial cell-death rate (see color bar) has four major zones: no stress zone (low salt – low antibiotics), antibiotic stress (low salts – high antibiotics), salinity stress (high salts – low antibiotics) and a zone of cross protection (high salts – high antibiotics). **B.** Examples of the trajectories of various initial antibiotic concentrations, with the resulting bacterial growth and death rates. Differing trajectories result in differing durations of cross-protection. In addition, antibiotic deactivation can lead to traversing the cross protection and returning to a no-stress zone after rewetting **C.** Schematic illustration of the mechanisms underlying the observed protection and their time of action during the various phases of a wet-dry cycle **D.** Dynamics of simulations, compared to observed results at the end of the long wet-dry cycle experiment and the corresponding ‘constantly wet’ experiments. **E.** Comparison of simulation results to observed results from a 17.5h wet-dry cycle and the equivalent ‘constantly wet’ experiment.

## Discussion

In this study, we demonstrated inherent protection of *E.coli* cells from two beta-lactam antibiotics, Ampicillin and Carbenicillin, under wet-dry cycles with MSW. We show that bacteria exposed to high concentrations of antibiotics – considerably higher than the MIC – showed significantly higher survival under a wet-dry cycle with prolonged ‘dry’ MSW conditions than they did under constantly wet conditions (Fig. 2).

Through a combination of experiments and computational modeling, we were able to point to four mechanisms operating in increased protection from beta-lactams under wet-dry cycles with MSW: (1) cross-protection due to high salt concentrations, (2) tolerance ‘by slow growth’, (3) deactivation of antibiotics by the physicochemical conditions associated with drying and MSW, and (4) tolerance ‘by lag’ (34) during rewetting. These mechanisms act at various phases of the wet-dry cycle (Fig. 4 C). Cross protection due to high salts and tolerance ‘by slow growth’ act in late drying and during the MSW phase, when antibiotic deactivation occurs. As deactivation is time dependent, the MSW duration modulates the accessible concentration of ‘active’ antibiotic after rewetting. Finally, the tolerance ‘by lag’ acts in the re-wetting phase. Our computational model enabled us to integrate the operation of these four mechanisms, and to analyze the effect of each mechanism, and its associated parameters, on the overall population response to antibiotics under wet-dry cycles.

It is difficult to assess the exact contribution of each of the first two mechanisms, i.e., cross protection due to high salts, and the tolerance ‘by slow growth’. This is partly due to the fact that activation of stress response mechanisms (e.g., from high salinity) that provides cross-protection from antibiotics, is intertwined with slow growth, which is known to lead to tolerance ‘by slow growth’ (34). While our experiments indicate that high salt concentrations are a major stress factor that induces cross protection (Fig. 3), other stresses associated with drying and MSW, such as altered pH and ROS, may be involved as well. Finally, we demonstrated that deactivation of the antibiotics was not enough to explain the observed protection. We note that we could not rule out the possibility that reduced bio-accessibility of antibiotics under MSW conditions played a role in this protection.

Identifying the molecular mechanism associated with the observed tolerance was not the focus of the current study, but rather aiming at a mechanistic understanding of the population-level response to antibiotics under wet-dry cycles and MSW. Possible molecular mechanisms facilitating reduced susceptibility associated with drying, rewetting, and MSW may include: expression of efflux pumps, general stress response and RpoS, oxidative stress response, reduced energy metabolism, SOS response, (p)ppGpp signaling, and toxin-antitoxin modules (39, 52–56).

Further research is required to identify the molecular mechanisms that facilitate the observed protection from beta-lactams under wet-dry cycles and MSW.

A surprising result was the significantly increased cross-protection from drying and MSW by the presence of lethal doses of antibiotics before drying. Although evidence for cross protection by antibiotics on other stresses, including osmotic stress in particular, has been reported, it was associated with resistant mutant strains, and not due to induced tolerance (57). Thus, substantial increased bacterial survival under MSW is, to the best of our knowledge, novel. Such prominent cross protection, showing even ~10-fold higher survival under wet-dry cycles if antibiotics are added (Fig. 2), may have significant implications for how we perceive antibiotics in the microbial world. For example, secretion of antibiotics may actually function to protect neighboring co-occurring species from drying, rather than killing them.

Our computational model was developed in order to achieve mechanistic understanding of bacterial response, at the population level, to beta-lactam antibiotics in wet-dry cycle with MSW. It thus aimed to validate that the proposed underlying mechanisms obtained from our experiments are sufficient to produce results that agree, at least qualitatively, with the experimental data. The model is relatively simple, and does not take into account the heterogeneity of individual cells within populations, nor the microscale heterogeneity of hydration and physicochemical conditions (2). Yet the model did enable us to highlight the mechanisms suggested as underlying the observed increased protection, and clarify their specific role in the response. The two-variable death rate function (Fig. 4A), was a key component of the model that enabled us to better understand the link between water level, salt and antibiotic concentration, and death rates. Accordingly, the trajectory on this death rate plane during a wet-dry cycle was illuminating. Additional or alternative mechanisms, or their details, are possible. For example, it was not clear whether the increased cross-protection from MSW stresses facilitated by the antibiotics depends upon the antibiotic concentration prior to drying, or the instantaneous one while under MSW (where some of the antibiotic activity decayed). In general, the model can be used to predict response to antibiotics and overall survival under various scenarios of hydration dynamics, initial antibiotic concentrations, and wet-dry cycle durations (Supp. Fig. S16). Further work is required to extend and refine the model, including additional experimental work to cover other antibiotics, additional variables, and a wider range of parameters.

In this work, we chose to focus on beta-lactams and the model bacterium *E. coli*. It is not clear whether the observed protection from beta-lactams is applicable to most bactericidal antibiotics. Cross-protection via increased expression of efflux pumps, for example, is a mechanism that has been demonstrated to reduce susceptibility to antibiotics of various families (46, 54), and depends upon chemical properties of the antibiotic (e.g., hydrophobicity). Tolerance by slow growth, however, is a general mechanism that has been demonstrated in various unrelated antibiotic families (e.g., beta-lactams and quinolones) (41–43). Also, as stability of various antimicrobial compounds may vary significantly, their degradation or deactivation under MSW may vary as well. Moreover, the present study did not explore antibiotic response of mature biofilms that may confirm tolerance due to matrix properties. Further research is thus required to assess these aspects, and to extend the study to other bacterial species.

This work changes how we understand bacterial response to a major class of antibiotics in several globally important microbial ecosystems, including the largest terrestrial microbial habitats: soil, rock surfaces, and plant root and leaf surfaces. It is also likely important to indoor environments, most of which are exposed to recurring wet-dry cycles with MSW, and of relevance to human skin microbiome. We demonstrate that in these vast, diverse microbial habitats, antibiotics act at a very different way than they do in test tubes or on lab plates. Under MSW conditions, for example, concentrations hundredfold higher than the clinical MIC are not very effective at killing bacteria. These findings have significant implications for how we understand antibiotics’ role in microbial warfare and competition, and the emergence of antibiotic resistance (58). They are also likely imperative for disinfection practices in indoor environments and for treating skin infections. To conclude, our results suggest that through their inherent properties, wet-dry cycles and the unique MSW environment may protect bacteria inhabiting vast diverse microbial habitats from major antibiotic groups.

## Methods

### Strains, media, chemicals, and growth conditions

*E. coli* Dh5α strain was transformed with a plasmid encoding a gene for blue fluorescent protein (BFP2) obtained from Mitja Remus-Emsermann (University of Canterbury Christchurch, NZ) via Addgene (plasmid #118492). In all experiments, a starter culture was grown overnight in M9 0.5X (M9 Minimal Salts Base 5x, Formedium, UK) supplemented with LB 1% (LB Lennox broth, Formedium, UK) and Glucose 20 mM, under agitation set at 220 rpm, at 37°C. Antibiotics used in the experiments were Ampicillin (Ampicillin sodium salt, Sigma) and Carbenicillin (Carbenicillin disodium, Formedium,UK).

### MIC Determination

50 μl of an overnight *E.coli DH5α* culture was transferred into 3 ml of fresh medium and incubated for an additional ~3h until OD_600_ reached a value of ~0.3 (1 cm optical path length). Cells were diluted into 96 microtiter plate (final OD_600_ = 0.005) set with a twofold serial dilution of Amp or Crb (10 serial dilutions from 100 μg/ml to 0.2 μg/ml) and a control without antibiotics (8 replicates for each dilution). Plates were incubated for 24h (at 32°C), then OD_600_ reads were taken in a plate reader (Synergy H1 Microplate Reader, BioTek Instruments, USA).

### Drying-rewetting experiments

#### Preparation

50 μl of *E.coli DH5α* overnight culture were transferred into 3 ml of fresh medium, and incubated for an additional ~3 h until OD_600_ reached a value of ~0.3 (1 cm optical path length). The OD_600_ = 0.3 culture was diluted to OD_600_ = 0.04 with fresh medium (as a control) or fresh medium supplemented with: (1) 5 μg/ml of Amp or Crb for the 2X MIC experiment (2) 50 μg/ml of Amp or Crb for the 20X MIC X20 experiment.

#### Wet-dry cycles and constantly wet conditions experimental setup

For the wet-dry cycle experiment, a 1 μl drop from each sample was deposited on the center of a well of a glass-bottom 24-well plate (24-well glass bottom plate #1.5 high-performance cover glass - Cellvis, USA). A total of six repeats (e.g., drops) was used for the 2X MIC and control experiments, and three repeats for the 20X MIC experiments. The 24-well plate was placed (with the plate lid off) inside a stage-top environmental control chamber (H301-K-FRAME, Okolab srl, Italy) pre equilibrated to 32°C, 70% RH. For the constantly wet experiment, a volume of 200 μl from each condition (two duplicates) was transferred to separate wells in a glass-bottom 96-well plate (96-well glass bottom plate #1.5 high-performance cover glass - Cellvis, USA), and placed, with the plate lid on, in an incubator set to 32°C.

#### Gradual rewetting protocol

At time t = 0, t = 3 and t = 14 h (times refers to the time that passed from deposition of the drops on the surface of the well), the empty cavities between the wells were filled with 300 μl of H_2_O, and the lid was placed on the plate and sealed with tape. The sealed plate was incubated for 90 min at 25°C (with RH raising to > 95% inside the plate). By the end of this step, the dried droplets (MSW) adsorbed water from the humidified air by condensation and deliquescence. At this point, 1 μl samples were pipetted out from the parallel ‘constantly wet’ experiment and deposited on a second 24-well plate, creating two identical sets of 24-well plates originating from (a) dry conditions (b) ‘constantly wet’ experiment. Next, 2 μl of fresh medium supplemented with 8 μM of propidium iodide (propidium iodide solution, Sigma) was added to the droplets of both wet-dry and constantly wet 24-well plates. The plates were left closed in the dark for 90 min, at 25°C. Finally, 4 μl of fresh medium was added to all rewetted droplets, and the plates were incubated for another 30 min under the same conditions.

#### Agarose pad overlay (CFU assay)

At the end of the rewetting procedure, 2 μl were pipetted out from each rewetted droplet and placed on the center of a precut round agarose pad (preparation of agarose pads described below). The remaining volume of the droplet was overlaid by a second agarose pad (round, 16mm diameter). Agarose pads were incubated for 6-12h in a humidified chamber at 25°C before imaging. 1.5% agarose pads were prepared by dissolving 1.5 g/l of agarose in M9 1X solution. The agarose-M9 solution was heated in a microwave until the agarose completely dissolved and a clear solution was obtained. The solution was cooled down to 55°C, supplemented with glucose 20 mM, and then poured into 10×10 cm petri dishes (~15 ml per dish). The agarose plates were left for 2h at room temperature, and then round disks of agarose pads were chopped using a 13 mm belt punch (ELORA, DE). Agarose pads’ overlay was done by picking an agarose disk using a flat spatula, placing the pad on a flat piston taken from a 10 ml syringe, and then carefully lowering the piston toward the rewetted droplet until contact between the pad and the glass surface was made. At this stage, the piston was pulled out while the agarose pad remained attached to the glass surface.

### Microscopy

24-well plates were mounted on a stage top without warming (at room temperature, 25°C) during image acquisition. Microscopic inspection and image acquisition were performed using an Eclipse Ti-E inverted microscope (Nikon) equipped with Plan Apo 20x/075 N.A. air objective and the Perfect Focus System for maintenance of focus. A LED light source (SOLA SE II, Lumencor) was used for fluorescence excitation. BFP fluorescence was excited with a 395/25 filter, and emission was collected with a T400lp dichroic mirror and a 460/50 filter. Propidium iodide fluorescence was excited with a 560/40 filter, and emission was collected with a T585lpxr dichroic mirror and a 630/75 filter (filters and dichroic mirror from Chroma, USA). A motorized encoded scanning stage (Märzhäuser Wetzlar, DE) was used to collect multiple stage positions. The entire surface of each drop was imaged at the end of the drying phase (BF and BFP channels) and after the addition of propidium iodide (BFP and RFP channels). In these images, 6×6 adjacent fields of view (with a 5% overlap) were scanned (3.83 × 3.83 mm). Agarose pads were imaged by scanning 20×20 adjacent fields of view (with a 5% overlap) (1.27 × 1.27 cm) in the BFP channel. Images were acquired with an SCMOS camera (ZYLA 19 4.2PLUS, Andor, Oxford Instruments, UK). NIS Elements 5.02 software was used for acquisition.

### Image Analysis

#### Image processing

Image processing and analyses was performed to quantify the amount of live (live/dead assay) and viable cells (CFU counts) in each experiment. Preprocessing of all images (i.e., background correction and threshold calibrations) was done in NIS Elements 5.02. For the live cell count within the drops, BFP and PI fluorescence channels of images were thresholded, and the resulting masks were united. Each cell in the unified mask was classified as a live cell if the fraction of its pixels marked with PI was lower than 10%. For the CFU count, BFP channel was thresholded to produce a mask. Only objects with an area larger than 12 μm^2^ (approximately 3 cells under our experimental conditions) were counted as CFUs. In some images, in particular in the ‘constantly wet’ experiments with high antibiotic concentrations, there were a few very large cells that had not divided. Manual inspection and correction was performed to avoid counting these large cells as CFUs.

#### Statistical analysis

The number of repetitions for each data point: 6 repeats for control experiments (low antibiotics), 6 repeats for low initial antibiotic concentrations (2X MIC x 2), and 3 repeats for high initial antibiotics concentrations (20X MIC).

### Estimating growth and lysis rates under various M9 concentrations

Growth and lysis rates were determined following Lee et al. (24) with slight modifications. 50 μl of an overnight *E.coli DH5α* culture was transferred into 3 ml of fresh medium and incubated for an additional ~3h until OD_600_ reached a value of ~0.3 (1cm optical path length). Cells were diluted into 96 microwell plates (final OD_600_ = 0.005) with a series of M9 concentrations of: 0.5X, 1X, 2.5X, 5X, 7.5X, 10X, and 15X (supplemented with LB 1% and glucose 20 mM). The plate was incubated in the plate reader for 4.5h (32°C, without shaking) and OD_600_ was monitored at intervals of 10 min. At time t ≈ 4.5h, the plate was pulled out, antibiotic (Amp or Crb) was added to the wells to give a final concertation of 50 μg/ml (equivalent to 20X MIC), and then transferred back to the plate reader for 20h with an OD_600_ scan every 10 min. Experiments were conducted in 3-5 replicates for each M9 concentration, and repeated three times with similar results. For the M9 10X and 15X experiments, bacteria were incubated for 24h in the tested M9 concentration to allow for cell acclimation prior to the addition of antibiotics.

#### Plate reader growth curve analyses

Plate reader readouts were analyzed as follows: First, the background value was subtracted from the OD values. Growth rates in absence of antibiotics were obtained by manual selection of a 2-3 hour duration in the log phase (typically between t = 5h to t = 7h) from the experiments without antibiotics (controls), and calculation of the slope. To estimate the lysis rates, we applied Gaussian weighing of adjacent readings (+10 and −10 minutes) to smooth the OD data, and then found the minimal rate of instantaneous change that occurs after antibiotic addition (i.e., at times > ~6h). The lysis rate was then estimated by substituting the growth rate without antibiotic, under the relevant control conditions from the minimal actual OD rate of change (See Fig. 3 and Supp. Fig. S11). The number of repetitions for each data point is five, except for three M9 concentrations (0, 5, and 7.5) in which there were three repeats for the control treatments, and four repeats for the antibiotic treatment.

### Computational model of antibiotic response

In this study, we developed an ODE model that describes antibiotic response of bacterial population under wet-dry cycles with an MSW phase. The model describes the dynamics of the following variables: number of live bacterial cells (*P*), water volume (*V*) [μl], and active antibiotic content by mass within the system (*A*) (normalized by [MIC × 1 μl]). The salt content by mass within the system (*S*_0_) (normalized by [Standard M9 concentration × 1 μl]) was constant throughout the simulation.

The external conditions are represented by the amount of imposed water volume at equilibrium *V_eq_(t)* [μl]. During the wet-dry-wet cycle, *V_eq_(t)* switches from *V_eq-wet_* to *V_eq-dry_* and *V_eq-dry_* to *V_eq-wet_* at times *t_W→D_* and *t_D→W_* respectively.

Under the above assumptions, we formulate the following ODE equation system:

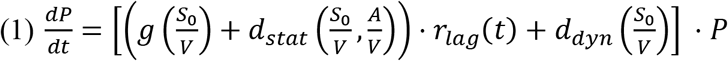

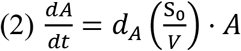

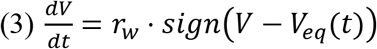

Note that 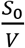 is the current salt concentration (*C_S_*) and *A*/*V* is the current antibiotic concentration (*C_A_*)

A more detailed description of the model is given in Section 1 of the Supporting Information.

### Measuring antibiotic stability

See Section 2 of the Supporting Information.

## Acknowledgements

We thank Jonathan Friedman, Nathalie Balaban, Yael Helman and Yitzhak Hadar for valuable comments and discussions. N. K. is supported by research grants from the James S. McDonnell Foundation (Studying Complex Systems Scholar Award, Grant #220020475) and from the Israel Science Foundation (ISF #1396/19).

## Author contribution

Y.B.M, M.G., T.O., and N.K. conceived the study. Y.B.M, and T.O. performed experiments. M.G. and T.O. performed image processing. M.G. and Y.B.M performed data analyses. M.G. conducted mathematical modeling and simulations. All authors discussed the results and contributed to the final manuscript. N.K. supervised the project. Y.B.M, M.G., T.O., and N.K. wrote the manuscript.

