## Supplemental Information for "Inherent protection of bacteria from beta-lactam antibiotics by wet-dry cycles with microscopic surface wetness"

### 1. Computational Model

In this study we have constructed an ODE model the can help explain the experimental results of wet-dry-wet cycle experiments mechanistically.

Several assumptions were made for the sake of simplicity: (a) bacterial growth is unbounded within the simulated duration, (b) antibiotic deactivation is caused by exposure to salts, and not by bacterial uptake, (c) water evaporates or condensates at a constant rate. Assumptions (a) and (b) are acceptable approximation at the conditions of our experimental system, where initial bacterial concentration is $2x10^4$ cells/μl. Assumption (c) is supported by our observations.

The model describes the dynamics of the following variables: The number of live bacterial cells ($P$), water volume ($V$) [μl], and active antibiotic content by mass within the system ($A$) (normalized by [MIC × 1μl]). The salt content by mass within the system ($S_0$) (normalized by [Standard M9 concentration × 1μl]) is constant throughout the simulation.

The external conditions are represented by the amount of imposed water volume at equilibrium $V_{eq}(t)$ [μl]. During the wet-dry-wet cycle, $V_{eq}(t)$ switches from $V_{eq-wet}$ to $V_{eq-dry}$ and from $V_{eq-dry}$ to $V_{eq-wet}$ at times $t_{W\to D}$ and $t_{D\to W}$ , respectively.

Under the above assumptions, we formulate the following ODE equation system:

$$(1)\ \frac{dP}{dt} = \left[\left(g\left(\frac{S_0}{V}\right) + d_{stat}\left(\frac{S_0}{V},\frac{A}{V}\right)\right)\cdot r_{lag}(t) + d_{dyn}\left(\frac{S_0}{V}\right)\right]\cdot P$$

$$(2)\ \frac{dA}{dt} = d_A\left(\frac{S_0}{V}\right)\cdot A$$

$$(3)\ \frac{dV}{dt} = r_w \cdot sign\big(V - V_{eq}(t)\big)$$

note that $\frac{S_0}{V}$ is the current salt concentration ($c_S$) and $A/V$ is the current antibiotic concentration ($c_A$).

Eq. 1: The terms of bacterial population ODE are defined as follows:

$g(c_S)$ denotes the growth rate of the bacterial population as a function of salt concentration:

$g(c_S) = \big(\mu(1 - sl(c_S, c_{S0}, K_S))\big)$, Where $sl(x, x_0, K) = \frac{1}{1+\exp(-K\cdot(log(x/x_0)))}$ is a sigmoid in logarithmic scale (see figure S11A)

$d_{stat}(c_S, \mathrm{c_A})$ denotes the death rate due to the instantaneous state of the system. It is a combination of sigmoid functions representing the response of cells to antibiotic and desiccation stress:

- At low salt concentrations, death as a response to antibiotics is modeled as a sigmoid function of the antibiotics concentration:
  $d^A_{lowS}(c_A) = d^A_{lowS} \cdot sl(c_A, c^A_{lowS}, K^A_{lowS})$ .
- At high salt concentrations, death as a response to antibiotics is modeled as a sigmoid function of the antibiotics concentration:
  $d^A_{highS}(c_A) = d^A_{highS} \cdot \left(1 - sl\left(c_A, c^A_{highS}, K^A_{highS}\right)\right)$ .

(Note that under high salt concentrations, death rates are higher with low antibiotic concentrations due to desiccation stress, and lower at high antibiotic concentrations due to cross-protection from high salts and desiccation.)

- At low antibiotic concentrations, death as a response to desiccation is modeled as a sigmoid function of the salt concentration:
  $d^S_{lowA}(c_S) = sl(c_S, c^S_{lowA}, K^S_{lowA})$ .

$d_{stat}(c_S, \mathrm{c_A})$ is a 2D function of both salt concentration and antibiotics concentration, created by combining $d^A_{lowS}(c_A)$, $d^A_{highS}(c_A)$ and $d^S_{lowA}(c_S)$:

$$d_{stat}(c_S, \mathrm{c_A}) = d^A_{lowS}(\mathrm{c_A}) \cdot \left(1 - d^S_{lowA}(c_S)\right) + d^A_{highS}(\mathrm{c_A}) \cdot d^S_{lowA}(c_S)$$

(see Supp. Fig. S12 B, C)

$r_{lag}(t)$ denotes the lag phase occurring after the MSW phase:

$$r_{lag}(t) = \begin{cases} 0 & t_{D\to W} \le t \le t_{D\to W} + t_{lag} \\ 1 & otherwise \end{cases}$$

$t_{lag}$ is determined by the length of the dry phase – modelled as a linear relation $t_{lag} = p_{lag} \cdot (t_{D\to W} - t_{D\to W})$

$d_{dyn}(c_S)$ denotes the death rate due to the transition between wet and dry states of the system:

$$d_{dyn}(c_S) = \frac{D^D}{2\pi \cdot \sigma} \cdot \exp\left(-\frac{log(x/C^D)^2}{2\sigma^2}\right),$$

which is a Gaussian at log scale, representing the reduction in cell numbers caused by transition from wet to dry conditions and vice-versa (See Supp. Fig. S13 C).

Eq. 2: Is the ODE of the antibiotic content.

$d_A(\mathrm{c_S}) = d^S_{deac} \cdot sl(c_S, c^S_{deac}, K^S_{deac})$ represents the rate of reduction in antibiotic activity caused by the physicochemical conditions associated with MSW, characterized by high salt concentrations in the solution (see Supp. Fig. S13 A, B).

Eq. 3: describes the evaporation and condensation process, when external conditions determine a different volume-at-equilibrium than the current water volume. The transition from dry to wet (or from wet to dry) conditions is represented by switching between $V_{eq-wet}$ and $V_{eq-dry}$ equilibrium values (or vice versa) during the simulation (see figure S1). The water volume approaches the current equilibrium at a constant rate $r_w$.

Model Parameters

Model parameters $\mu, r_w$, $d^A_{lowS}$, $c^A_{lowS}$, $K^A_{lowS}, d^S_{lowA}$, $c^S_{lowA}$, $K^S_{lowA}$, $d^A_{highS}$, $d^S_{deac}$, $c^S_{deac}$, $K^S_{deac}, D^D, \sigma, C^D$ were calibrated by experimental data as shown in Supp. Figs. S12 and S13.

Model parameters $c^A_{highS}$, $K^A_{highS}$, $t_{lag}, p_{lag}$ were chosen to fit the experimental values of bacterial population at the end of a wet-dry cycle.

The model was simulated in MATLAB using the standard ODE45 solver

Model parameter values:

| Name | Value | Units |
|---|---|---|
| $P_0$ | 22,000 | Cells |
| $A_0$ | 0.01, 2 or 20 | ×[MIC] |
| $V_{eq-wet}$ | 1 | $\mu l$ |
| $V_{eq-dry}$ | 0.01 | $\mu l$ |
| μ | 0.19 | $h^{-1}$ |
| $c_{S0}$ | 7.5 | ×[Standard M9 concentration] |
| $K_S$ | 20 | |
| $r_w$ | 0.5 | $h^{-1}$ |
| $d^S_{lowA}$ | 0.36 | $h^{-1}$ |
| $c^S_{lowA}$ | 7.5 | ×[Standard M9 concentration] |
| $K^S_{lowA}$ | | |
| $D^D$ | 5 | $h^{-1}$ |
| $C^D$ | 0.1 | ×[Standard M9 concentration] |
| $\sigma$ | 1 | |
| $t_{lag}$ | 3 | $h$ |
| $p_{lag}$ | 0.25 | |
| Ampicillin | | |
| $d^A_{lowS}$ | 1.19 | $h^{-1}$ |
| $c^A_{lowS}$ | 2.1 | ×[MIC] |
| $K^A_{lowS}$ | 10 | |
| $d^A_{highS}$ | 0.36 | $h^{-1}$ |
| $c^A_{highS}$ | 40 | ×[MIC] |
| $K^A_{highS}$ | 10 | |

| | | |
|---|---|---|
| $d_{deac}^{S}$ | 0.17 | $h^{-1}$ |
| $c_{deac}^{S}$ | 12.5 | ×[Standard M9 concentration] |
| $K_{deac}^{S}$ | 1000 | |
| Carbenicillin | | |
| $d_{lowS}^{A}$ | 0.79 | $h^{-1}$ |
| $c_{lowS}^{A}$ | 1.5 | ×[MIC] |
| $K_{lowS}^{A}$ | 10 | |
| $d_{highS}^{A}$ | 0.36 | $h^{-1}$ |
| $c_{highS}^{A}$ | 80 | ×[MIC] |
| $K_{highS}^{A}$ | 10 | |
| $d_{deac}^{S}$ | 0.083 | $h^{-1}$ |
| $c_{deac}^{S}$ | 12.5 | ×[Standard M9 concentration] |
| $K_{deac}^{S}$ | 1000 | |

### 2. Measuring antibiotic stability

Bacteria were used as a bio-reporter for antibiotic activity. Antibiotic deactivation was estimated as percentage of the original activity over time (at 1 h, 4 h, 8 h and 24 h), under drying and MSW conditions, or at various M9 concentrations.

*Antibiotic stability in MSW*: for each tested time point, 15 1-μl droplets of M9 0.5X supplemented with 1000 μg/ml of antibiotic (Amp or Crb), were deposited on the surface of two wells of a glass bottom 24-well plate (a total of 30 droplets). Droplets were dried under the same conditions described in Methods (*Wet-dry cycles and constantly wet conditions experimental setup*). At the end of the incubation period (1, 4, 8 or 24 h), each well was washed with 150 μl of fresh medium (without antibiotics), which results in the re-suspension of the aged antibiotic to a final concentration of 100 μg/ml. The volume of the two replicated wells was mixed and used as a starting stock for a serial dilution MIC assay (see above, *MIC Determination*). The MIC experiment was done in two parallel plates: one with aged antibiotics and the other with fresh antibiotic. The activity of the aged antibiotic (%) was determined by comparing between the measurements of final bacterial $OD_{600}$ after 24 hours of incubation.

*Antibiotic stability in varying M9 concentrations*: In this experiment antibiotics were incubated in bulk conditions in four M9 concentrations: 1X, 5X, 10X and 15X for a duration of 1, 4, 8 and 24 h (at $32^{0}$C). The initial antibiotic concentration in each experiment was adjusted such that after dilution of the concentrated M9 to M9 1X, the antibiotic concentration will be 50 μg/ml (assuming 100% stability). At each time point, the modified media was diluted with the required M9 concentration or DDW to get M9 2.5X and antibiotic concentration of 50 μg/ml. Antibiotic activity was determined by comparing aged and fresh antibiotic as described above (*Antibiotic stability in MSW*).

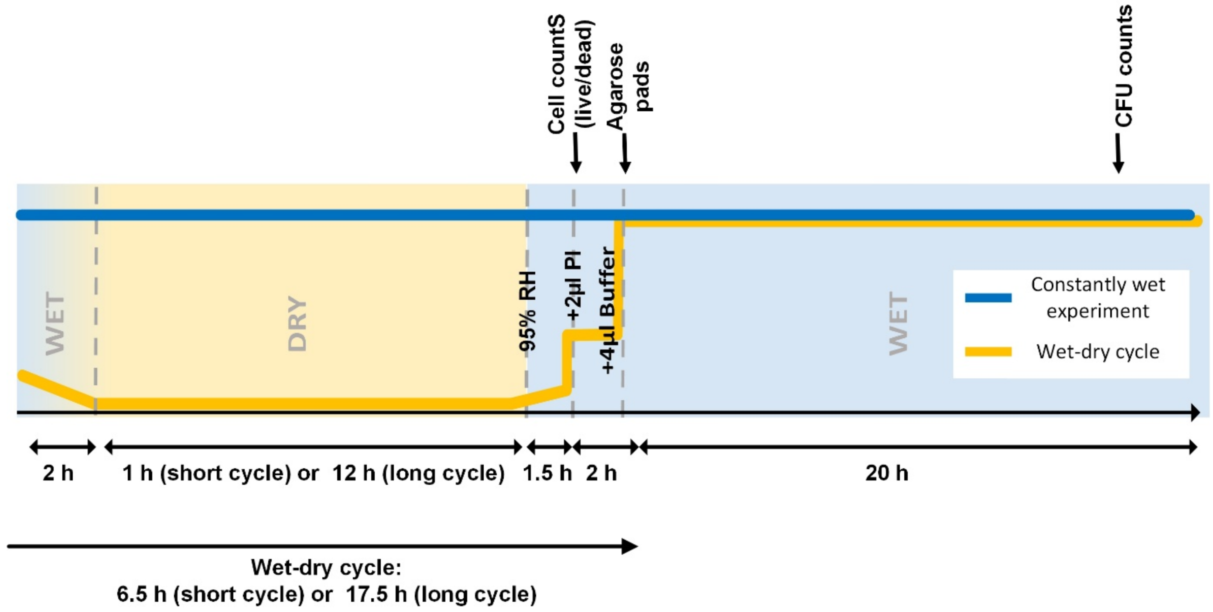


**Supp. Fig. S1. Wet-dry cycle and MSW experimental scheme.** In the wet-dry cycle with MSW experiments, 1µl drops were loaded and dried under constant temperature and humidity ($32^0$C and 70% RH, respectively). After 1h (short cycle) or 12h (long cycle) under MSW conditions, rewetting was done by first increasing RH to >95% (for 1.5h) followed by addition of 2 µl of medium with Propidium iodide. At this point Live/dead assay was conducted. Then additional 4 µl medium was added and agarose pads were laid over for a CFU counts (following further incubation of ~20h). According to this scheme the durations of the short and long wet-dry cycles in our experiments were 6.5h and 17.5h, respectively. See details in Methods.

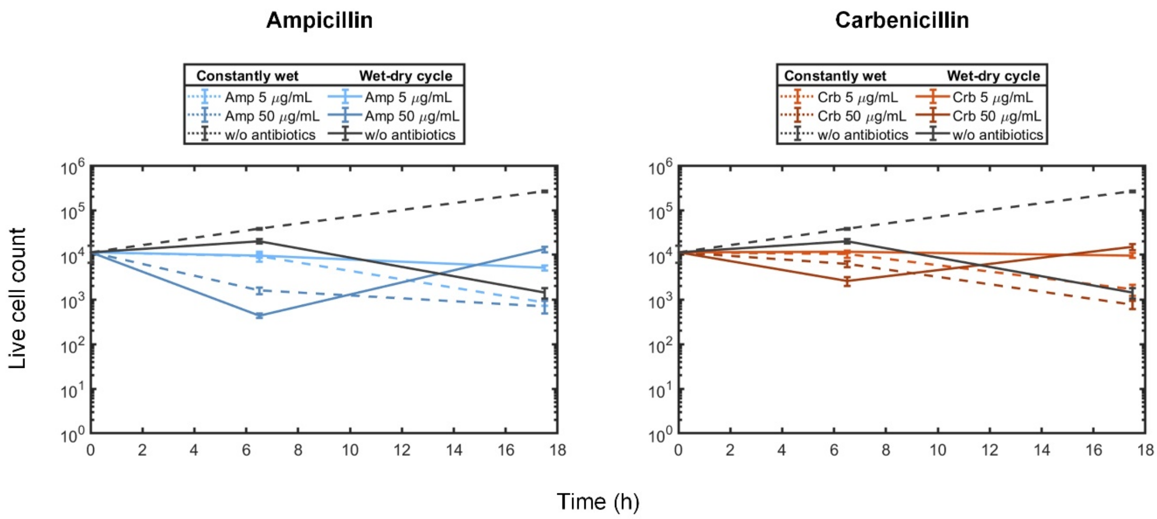


**Supp. Fig. S2. Antibiotic response to beta-lactams, live/dead assay results. A.** Bacterial response to Amp under short and long wet-dry cycle, and in constantly wet (water-saturated) conditions. Mean ± SE (number of live bacteria per initial 1 µl drop) are at times 0, 6.5 h and 17.5 h. Amp was added at concentrations of 5 µg/ml and 50 µg/ml; no antibiotic was added in controls. Live/dead quantification assay was done as described in Methods. **B.** same as (A) but with Crb.

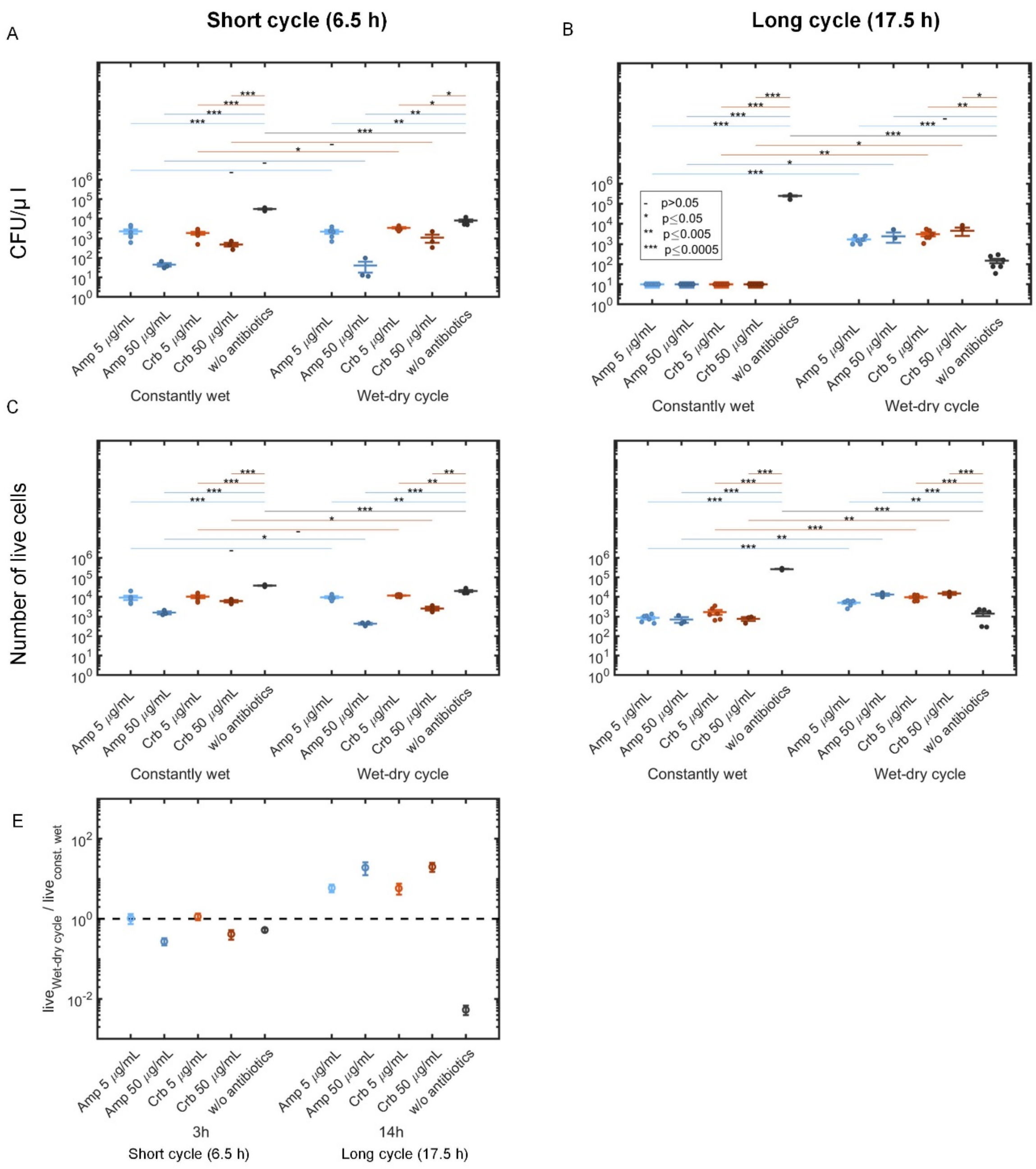


**Supp. Fig. S3. The number of live bacterial cells or CFUs at the end of wet-dry cycles and constantly wet experiments.** Number of CFU at the end of a short 6.5h (**A**) and long 17.5h (**B**) wet-dry cycles. Number of live cells (live/dead assays) in a short 6.5h (**C**) and long 17.5h (**D**) cycles. Individual points represent replications (individual drops). Horizontal lines and symbols indicate the significance of two-sample t-test between relevant pairs of experiments. (**E**) Ratios of live cells counts in wet-dry cycles with MSW and the corresponding constantly wet experiments (similar to Fig. 2C main text, but for live cell counts instead of CFU counts). Note the ~10-fold higher live cell counts in the long cycle experiment in comparison to constantly wet conditions.

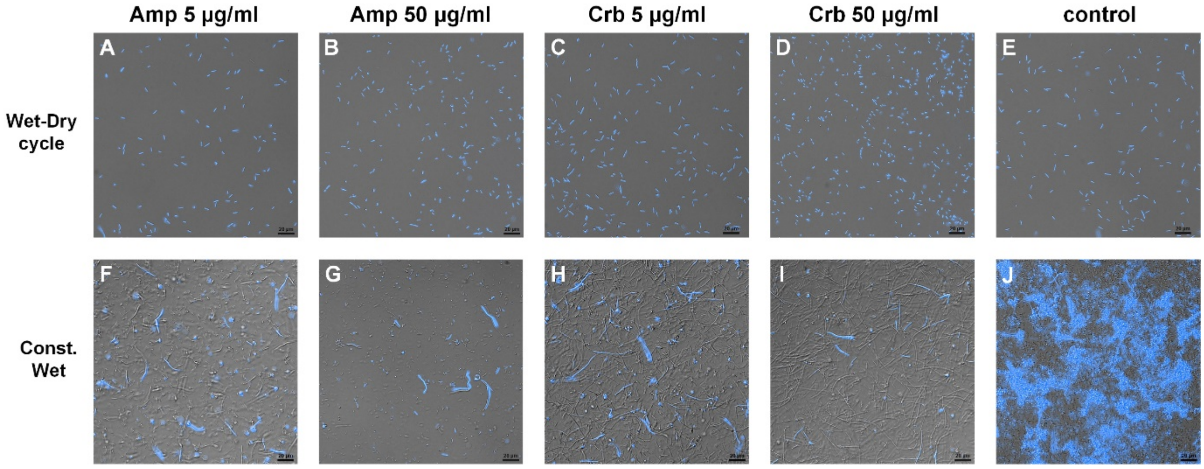


**Supp. Fig. S4. Representative sections of the surface imaged 14 h after drying experiment initiation.** The upper row (A-E) captures a surface section at the end of an MSW phase under the tested treatments. The lower row (F-J) capture a surface section in the constantly wet experiment under the tested treatments. In the constantly wet experiment with antibiotics (F-I), many of the cell changed their morphology and became longer, a known response to bet-lactams. In addition, the majority of cells do not express BFP indicating these cells are not alive. Images show a 0.25 × 0.25 mm section of the surface.

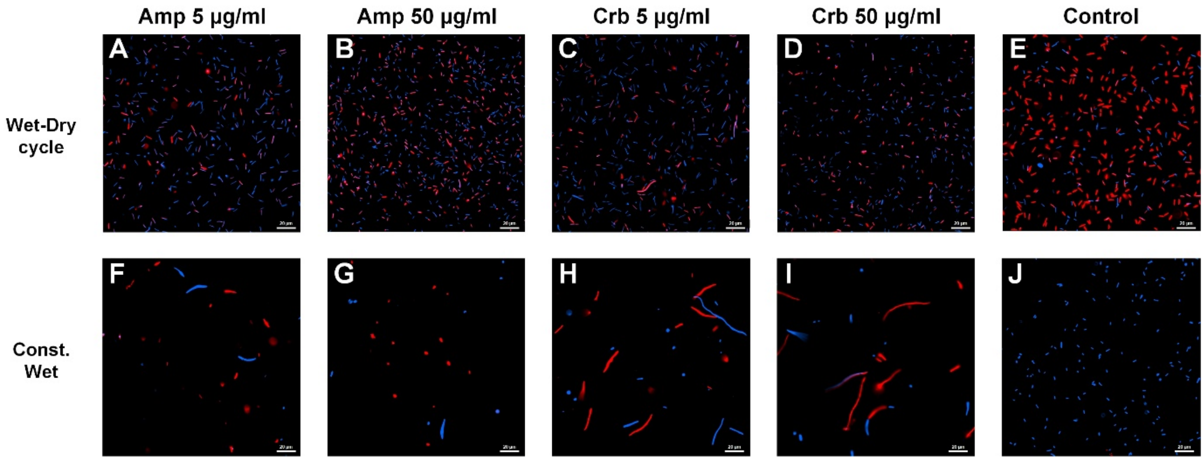


**Supp. Fig. S5. Representative sections of the surface imaged during the rewetting of the long wet-dry cycles experiment (at t=15.5 h), and the corresponding constantly wet experiments, both with propidium iodide (PI) staining**. The upper row (A-E) capture surface section during the rewetting (after 12 h in MSW conditions). Under most treatments, less than ~40% of the cells were stained with PI (e.g., red cells) which indicates that most cells are intact. In the control experiment (without antibiotics, panel E) most of the cells are stained with PI which implies that survival rate was lower than in the treated samples. The lower row (F-J) capture surface sections from a 1 µl drop sampled from the constantly wet experiment. Under all treatments (except control, panel J) the concentration of fluorescent cells (blue and red) is lower than in the equivalent treatment in the corresponding wet-dry cycle experiment. This may be explained by the high rate of burst cells that neither express BFP or stained with PI. The BFP expressing cells may look a live but not necessarily viable (not forming CFUs). The cells from the control sample (panel J) was 10-fold diluted for ease of image presentation. These cells express BFP and are not stained with PI which indicates that cells are alive. Images show a 0.26 × 0.26 mm section of the surface.

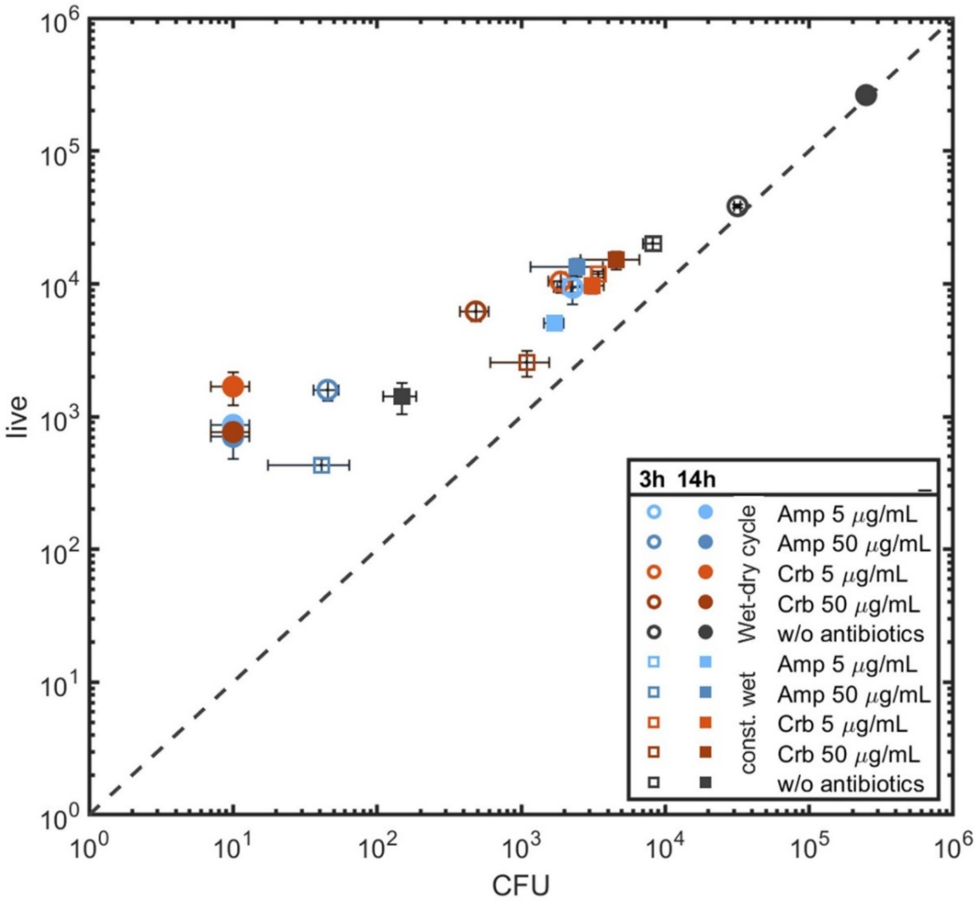


**Supp. Fig. S6. Correlations between live/dead assays and CFU counts.** 3h data represent short cycle; 14h represent long cycle. Pearson's correlation coefficient: 0.9979 (p-value: 8e-23). F-test: 0 (p-value: 0.85).

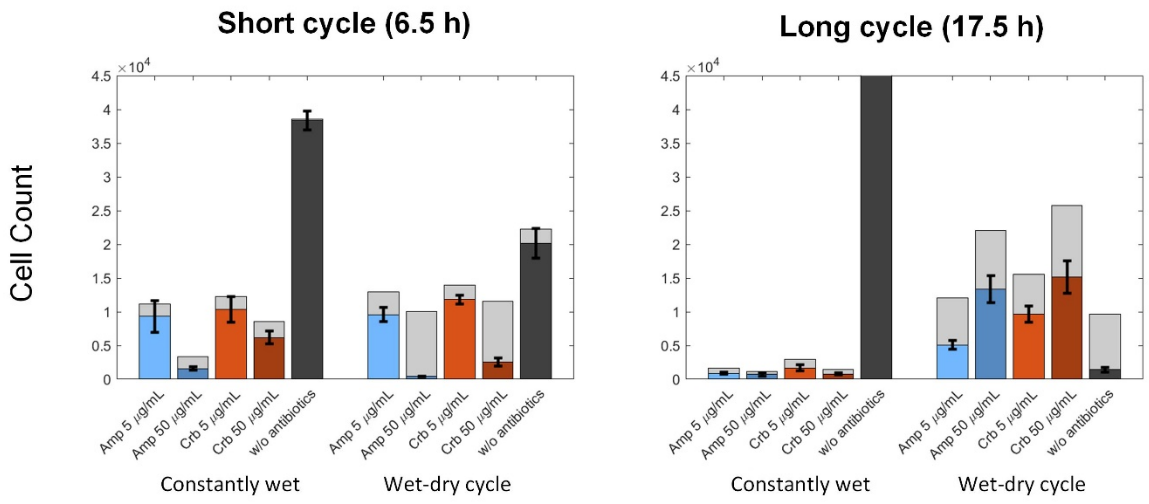


**Supp. Fig. S7. Total live and dead cell number at the end of wet-dry experiments (based on live/dead assays)**. Proportions of live (colorful) and dead cells (light grey) in the short (left) and long (right) wet-dry cycle experiments. Number of cells is per drop (1µl initial volume). Note the low live-to-dead cells ratios in the high antibiotic concentrations in the short cycles, in comparisons to the same ratios in the long cycle experiments.

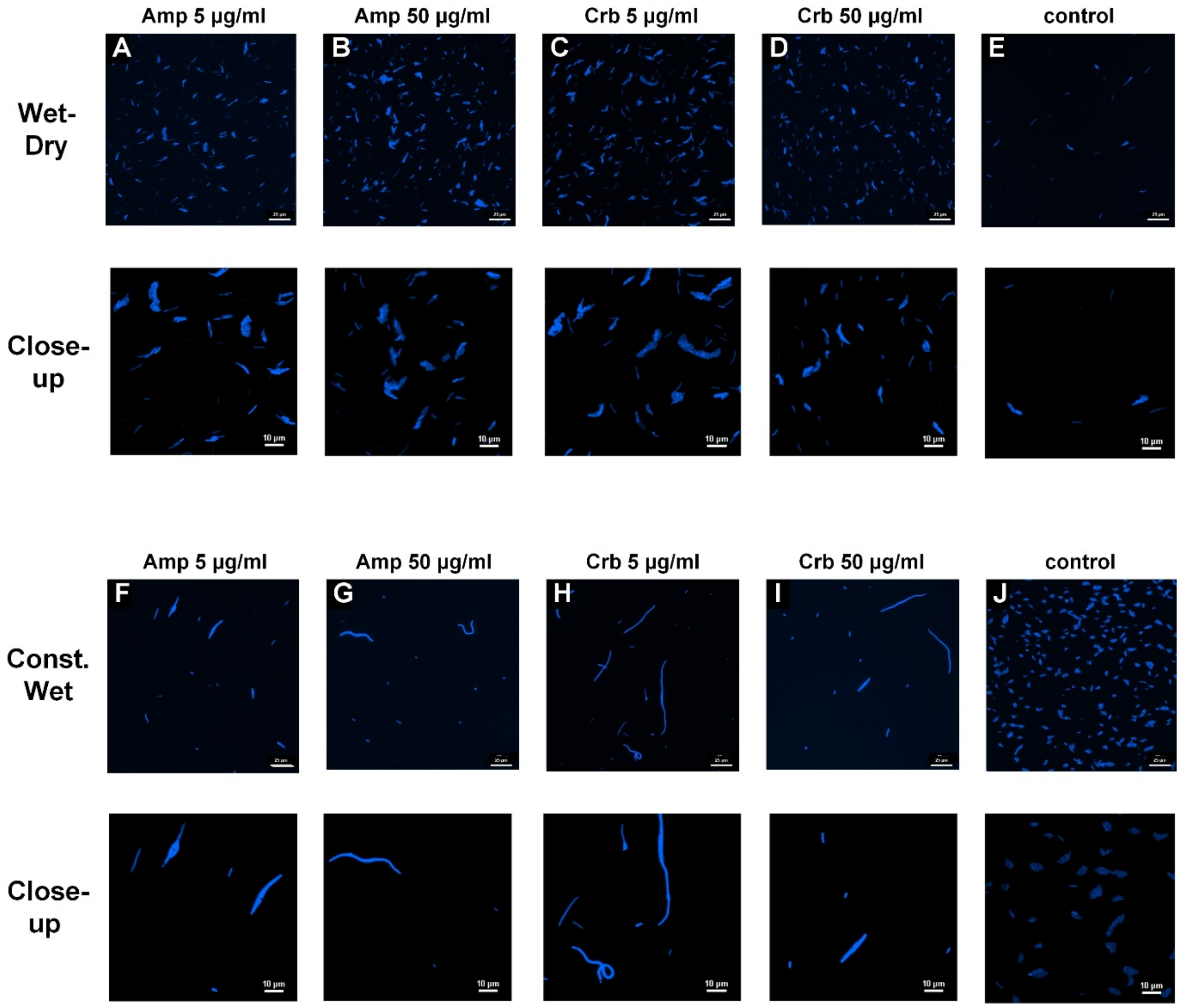


**Supp. Fig. S8. Representative sections from agarose pads from wet-dry and constantly wet experiments (CFU assays).** The first row (A-E) shows sections on the agarose, 16 h after it was overlaid on the bottom surface of rewetted 1 µl drop (long wet-dry experiment). Micro-colonies are observed in all antibiotic-treated samples (panels A-D) under wet-dry cycles. In the control experiment (e.g., without antibiotics, panel E), the number of micro-colonies was significantly lower than in the antibiotic treatment samples, indicating on higher cell viability in the presence of antibiotics. The third row (F-J) capture sections of the agarose pad taken from the corresponding constantly wet experiment. With antibiotics, micro-colonies are rarely observed, only extremely elongated cells expressing BFP were commonly noticed. In contrast, in the control experiment without antibiotics (panel J) the agarose was covered with micro-colonies (the sample was diluted by 10 folds to allow CFU count). The images in the second and the fourth rows are close-up images of the first and the third rows, respectively. Images show a 0.26 × 0.26 mm section of the surface.

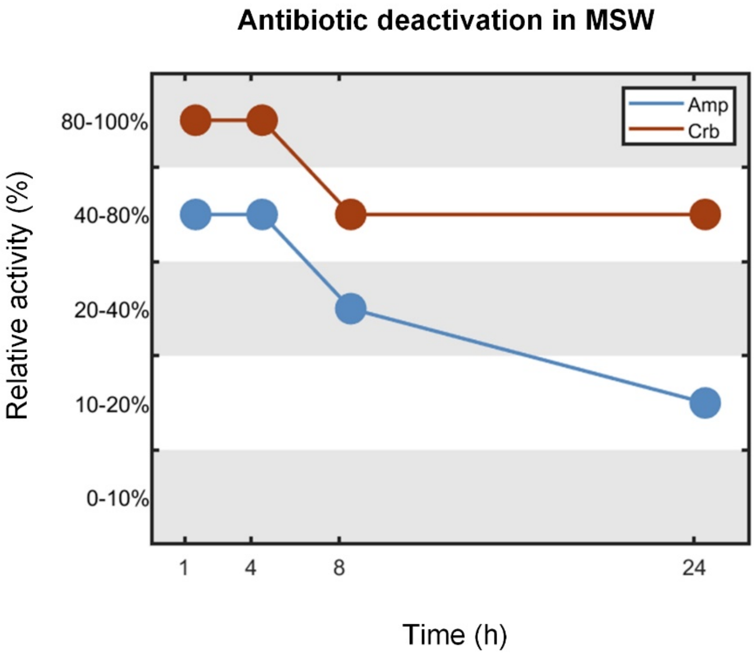


**Fig. S9. Antibiotic deactivation under drying and MSW conditions.** Amp and Crb stability was estimated as described in Methods. Amp is deactivated faster than Crb, with a reduction of more than 50% activity after 8 hours under MSW conditions. Amp deactivation was more pronounced: it showed 10-20% remaining activity after 24 h in MSW, in comparison to 40-80% remaining activity of Crb (Fig 3, Methods). This deactivation of antibiotics is explained, at least in part, by the high salt concentrations (Supp. Fig. S10).

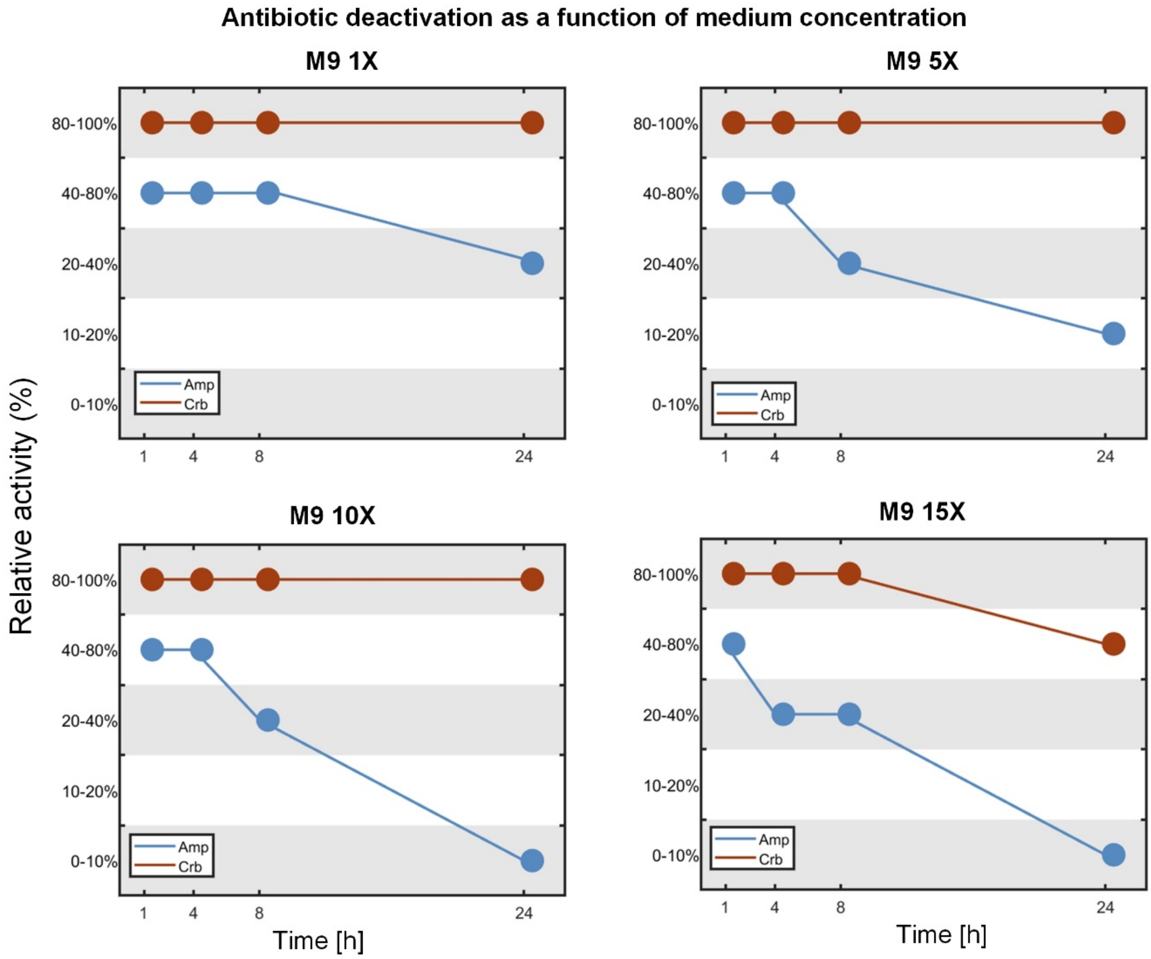


**Supp. Fig. S10. Antibiotic stability as a function of medium concentrations.** Stability of 50 µg/ml Amp (blue) and Crb (orange) in M9 1X, M9 5X, M9 10X and M9 15X.

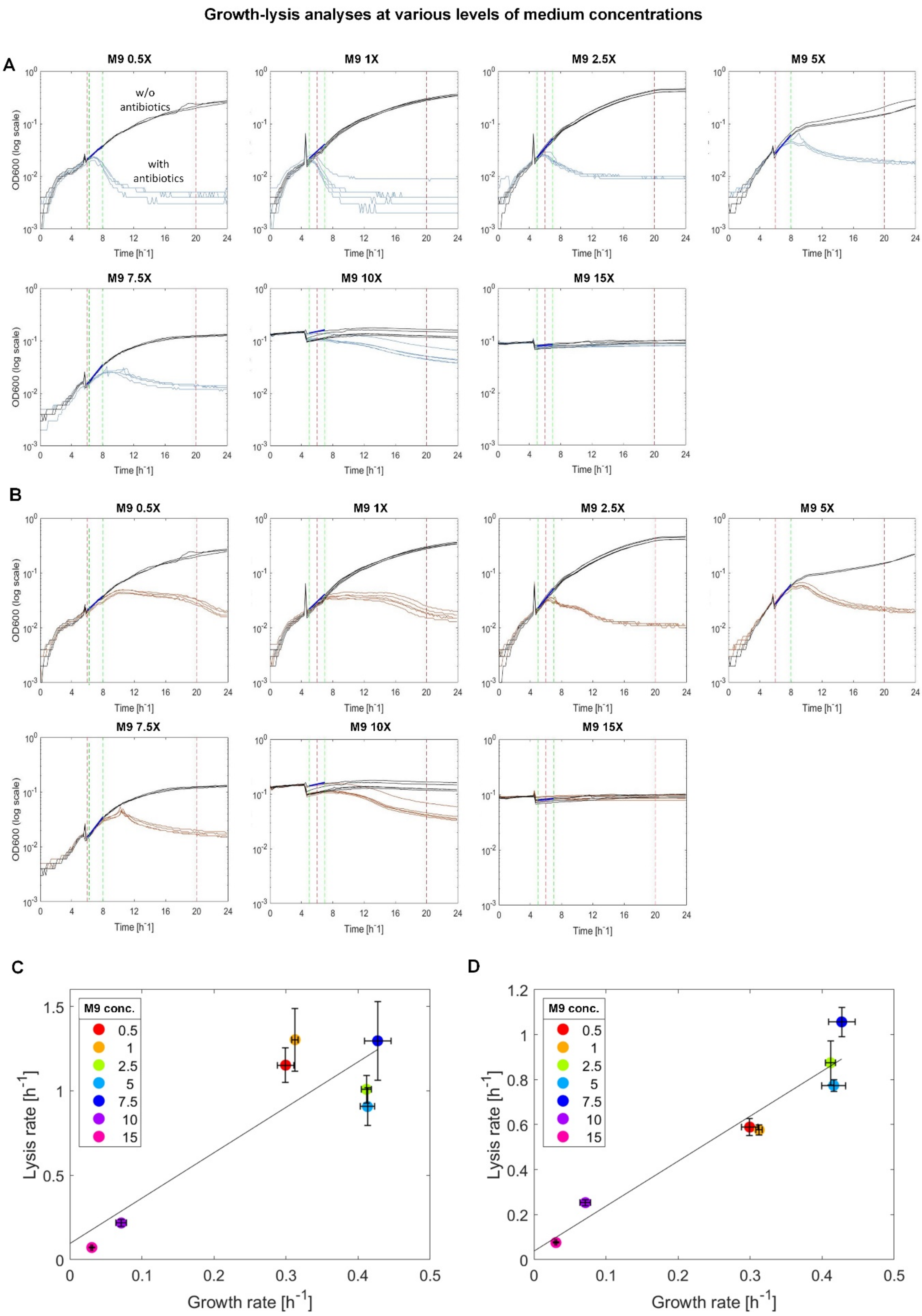


**Supp. Fig. S11. Bacterial growth curves and analysis. A.** Growth curves in increasing M9 concentrations without antibiotics (black lines) and with 50 µg/ml of Amp (blue lines). Antibiotic was added 4-6 h after inoculation. The vertical dashed green lines represent time range used to compute growth rates. The vertical dashed red lines represent time range used to compute minimal change rate in OD. **B.** Similar to (A) but with 50 µg/ml of Crb (orange lines). **C-D.** correlation between lysis and growth rates in increasing M9 concentrations (mean ± SE). Line equation for Ampicillin is 1.7x + 0.089 (R^2 = 0.84) and for Crb it is y = 1.5x + 0.044 (R^2 = 0.98). Pearson's correlation coefficient for Amp is r=0.84 (p-val 1.9e-2) and for Crb it is r=0.98 (p-val 5.7e-5). F-test for Amp is 0 (p-val 9.8e-2) and for Crb F-test is 0 (p-val 0.36).

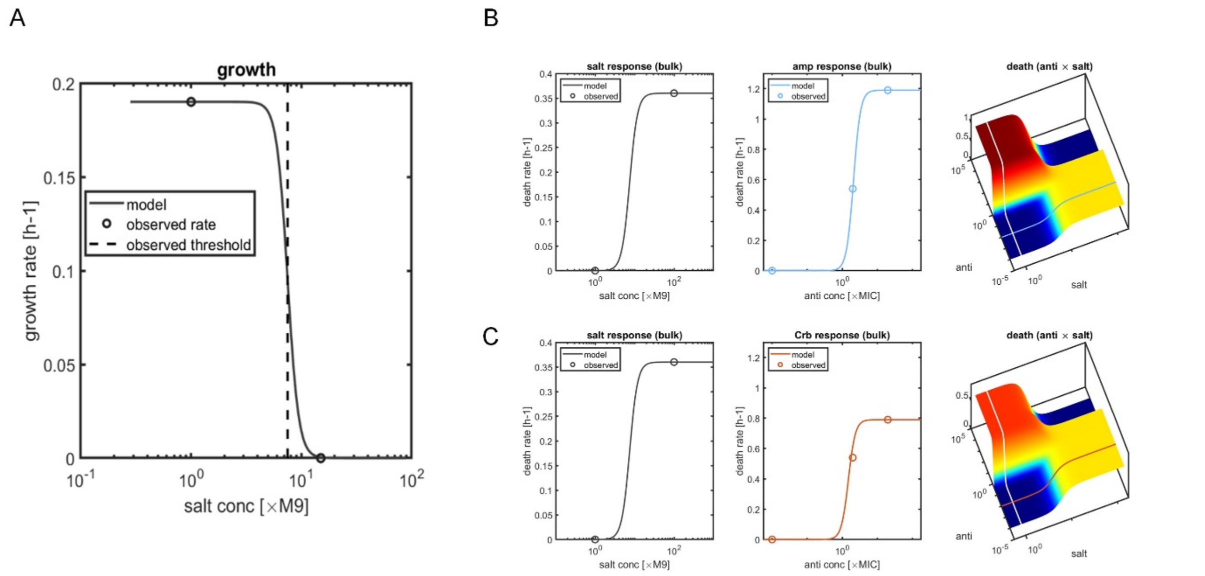


**Supp. Fig. S12. Calibration of model parameters**. **A.** Growth rate as a function of salt concentrations: Maximal growth rate was calculated based on comparing CFU counts of the bulk experiments without antibiotics at 6.5 h and 17.5 h (see Fig. 2 A-B). The critical salinity value for the growth rate function was determined based on the constantly wet experiments under constant salinities (see Fig. 3 in main text) **B-C.** Death rates as a function of Ampicillin (panel B) and Carbenicillin (panel C) concentrations and salt concentrations: Left: Death as a function of salinity without antibiotics was calculated by CFU counts of the wet-dry cycle and constantly wet experiments without antibiotics (see Fig. 2 A-B main text). Middle: Death as a function of antibiotics at low salinity was calculated by comparing CFU counts of the constantly wet experiments with high, low or without antibiotics at 6.5 h. Right: The two-variable (salt concentration and antibiotics concentration) death function is a combination of both functions, with the added assumption of cross protection at high salt concentration and high antibiotics concentration.

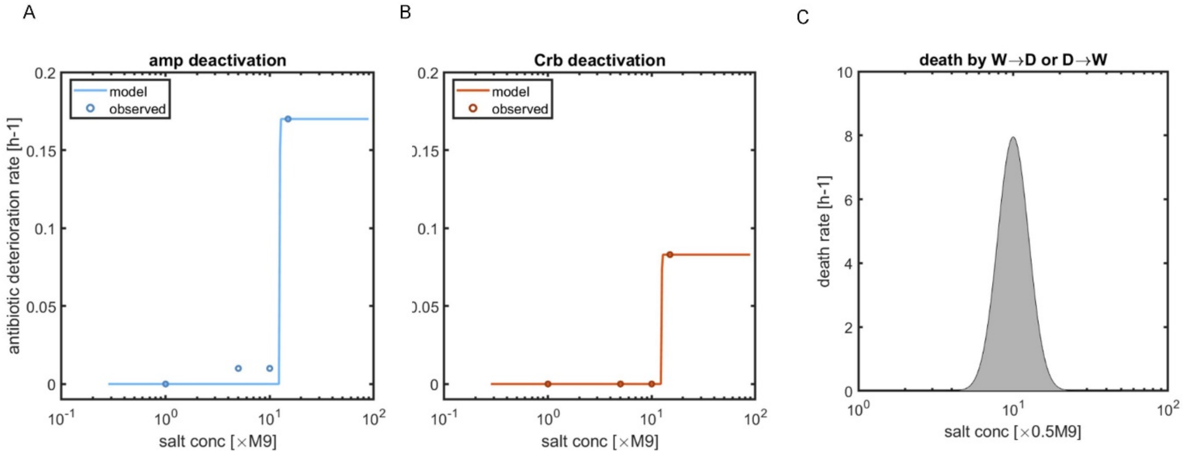


**Supp. Fig. S13. Calibration of model parameters 2. A-B.** Ampicillin (panel A) and Carbenicillin (panel B) deactivation rate. The rate was approximated as a sigmoid function of salinity, with maximal values drawn from deactivation rates for experiments at 24 h (see Fig. S9 in main text). **C.** Death resulting from the rapid transitions between wet (water saturated) and MSW states (drying) and back to wet conditions (rewetting) is modeled here, as increased death at intermediate salt concentration values. The overall mortality induced by both transitions was chosen to fit short- and long wet-dry cycle experiments with no antibiotics.

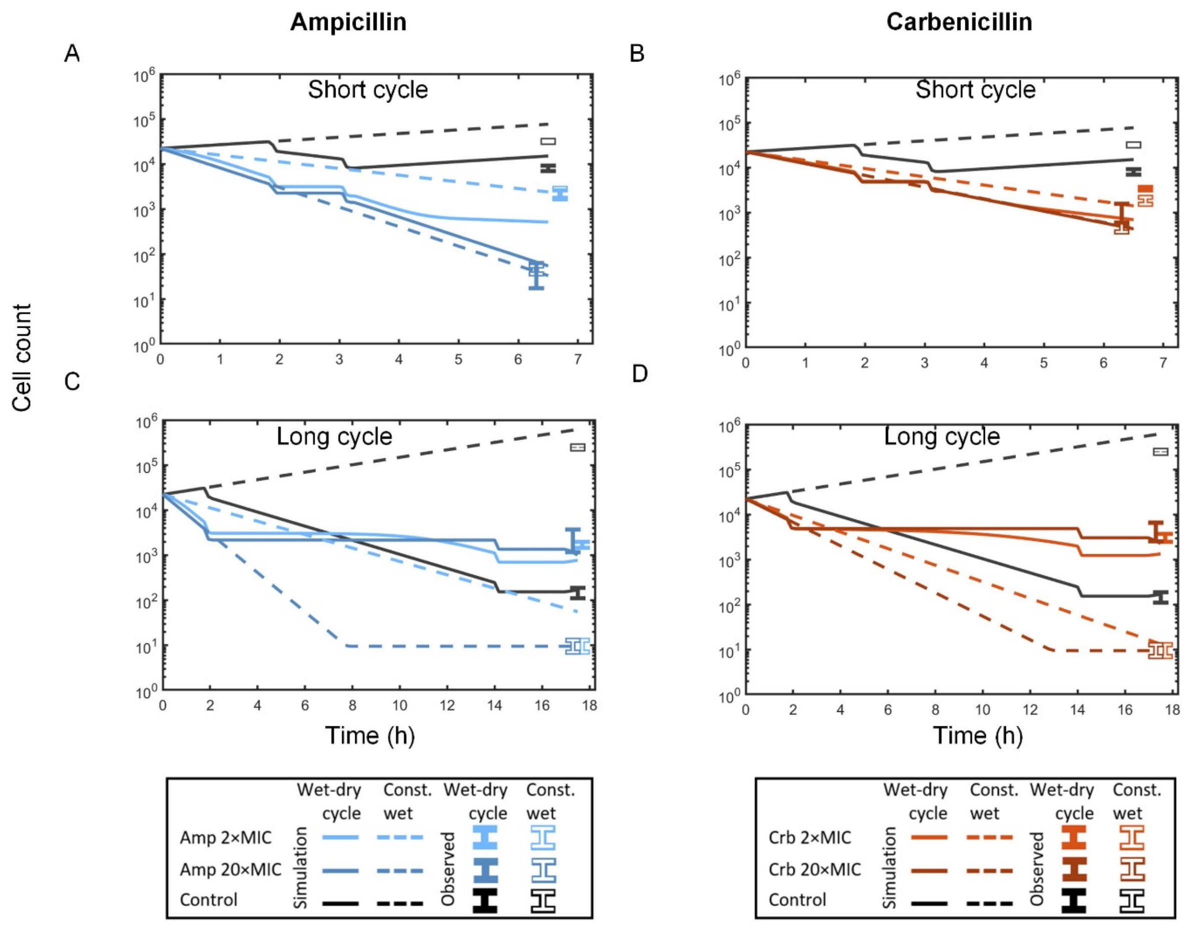


**Supp. Fig. S14. Simulations of population dynamics in short and long wet-dry cycles.** Simulations results are compared to observed experimental results for Ampicillin (left) and Carbenicillin (right), at the end of the 6.5 h (left) and 17.5 h (right) wet-dry cycle experiment and the corresponding constantly wet experiments. Experimental results (mean ± SE) are based on CFU counts (in CFU/μl units).

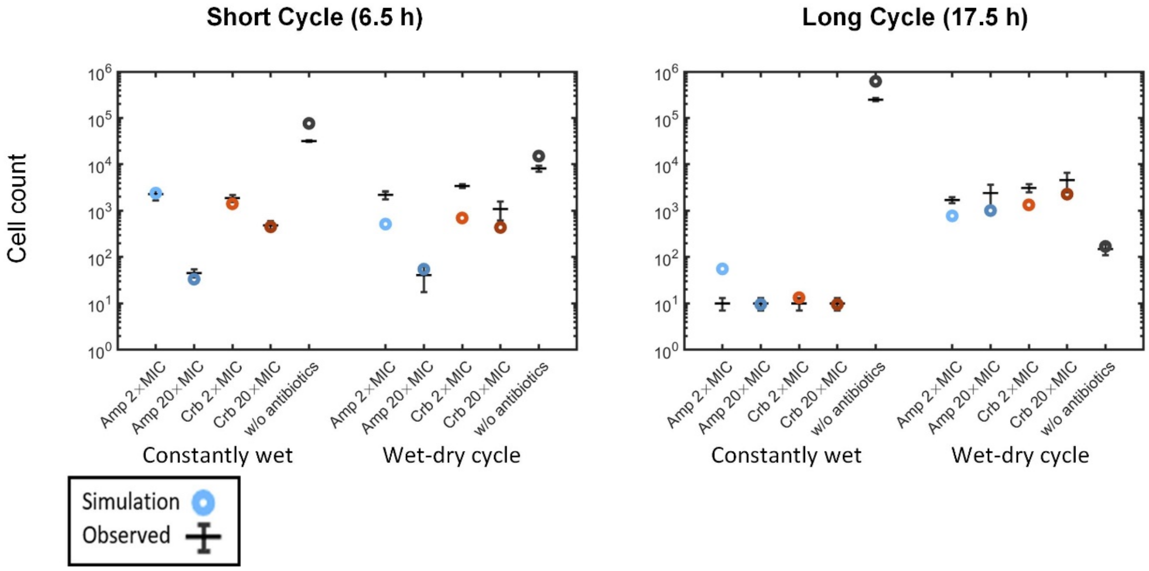


**Supp. Fig. S15. Comparison between experimental results and model simulations.** Comparison of simulation results to observed result from a short (left) and long (right, shown in Fig. 5D) wet-dry cycles and the equivalent constantly wet experiments. Experimental results (mean ± SE) are based on CFU counts (in CFU/µl units).

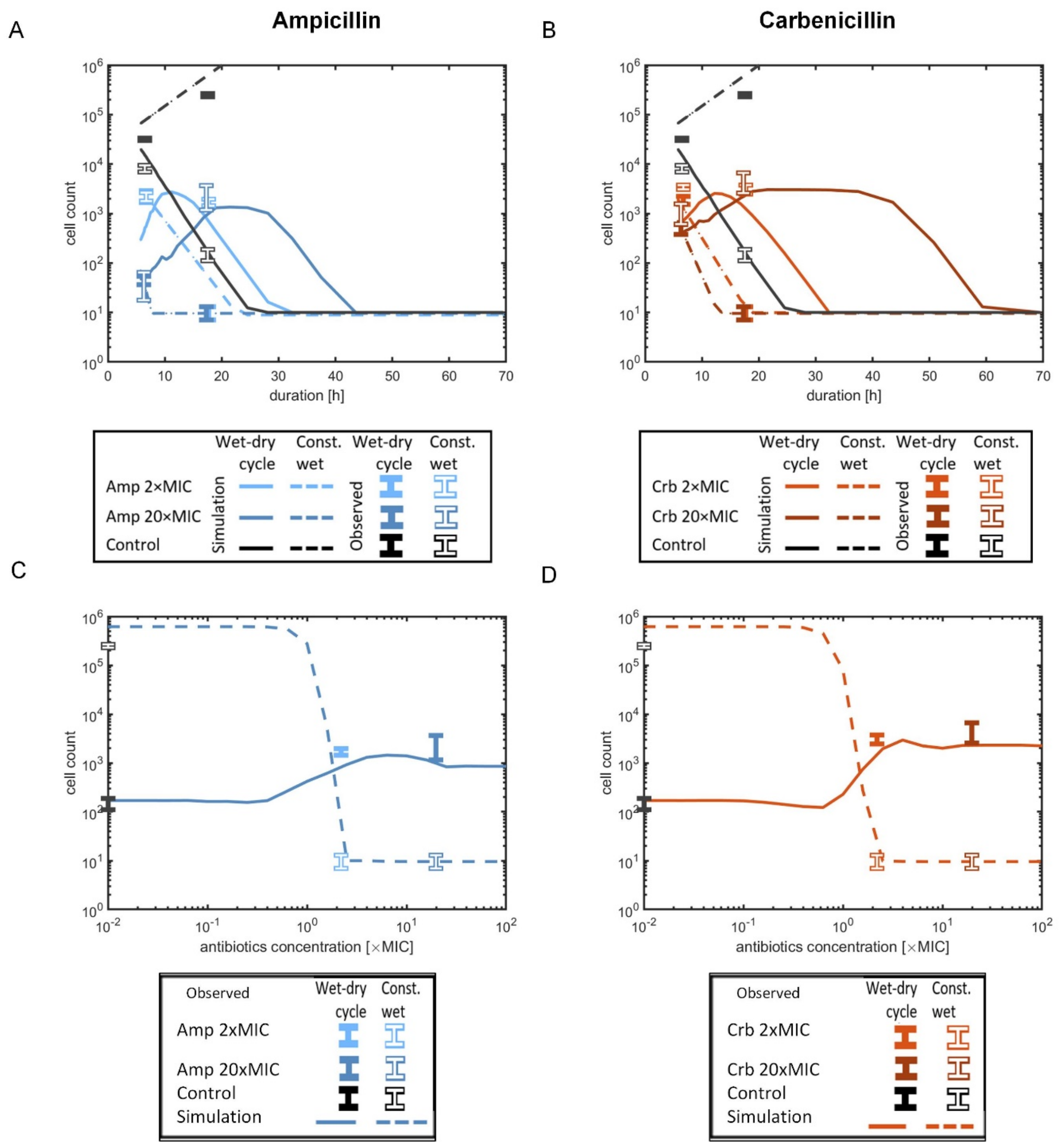


**Supp. Fig. S16. Projection of model results. A-B.** Population size as a function of wet-dry cycle length for Ampicillin (panel A) and Carbenicillin (panel B). **C,D:** Population size as a function of initial antibiotic concentration for Ampicillin (panel C) and Carbenicillin (panel D). Experimental results (mean ± SE) are based on CFU counts (in CFU/µl units).
